# Complementary attentional mechanisms for the resolution of representational ambiguity in the human brain

**DOI:** 10.1101/2025.09.22.677741

**Authors:** Clayton Hickey, David Acunzo, Damiano Grignolio

## Abstract

When we view cluttered environments containing multiple objects, the neural code for individual items can become ambiguous. This reflects a capacity limit in spatially-tuned visual neurons: when multiple objects fall within the cell receptive field, output cannot be attributed to a single object. Vision models from primate electrophysiology propose this is resolved by attention, which biases competition between object representations. However, evidence in humans is sparse and inconclusive, and reliance on single-cell data limits insight into underlying mechanisms and neurophysiological scope. Here, we use a novel multivariate approach to the analysis of human EEG and concurrent EEG/MRI to address this. First, we test whether attention is recruited by representational ambiguity. Second, we identify the mechanisms that act on representations of attended and unattended objects to resolve ambiguity. Finally, we characterize the millisecond timing and whole-brain action of these mechanisms to identify pervasive effects in semantic and executive brain networks.

## Introduction

The human visual system is remarkably efficient, yet fundamentally constrained: we are bombarded with more visual information than we can effectively process or act on. A class of theoretical models, centred on *biased competition theory* (Desimone & Duncan, 1995) and *ambiguity resolution theory* (Luck, Girelli, McDermott, & Ford, 1997; see also Usher & Niebur, 1996; Tsotsos, 1995; Deco & Rolls, 2004), suggest this capacity limit is resolved through attentional selection. This idea is grounded in the observation that early visual cells are spatial in nature: they respond to stimuli that fall within retinotopic receptive fields (RFs). When two stimuli fall within the RF of one cell, and that cell is sensitive to both, neural output can no longer be attributed to one item. To resolve this, attention is thought to bias the competitive interaction of object representations so cells unambiguously represent the attended object.

Electrophysiological studies in non-human primates have provided strong evidence for this idea. When two stimuli fall within the receptive field of a single extrastriate neuron, the firing rate is not the sum of responses to each object individually. Critically, attending to one of the objects shifts the response toward what would be observed if it were presented alone (Moran & Desimone, 1985; Chelazzi, Miller, Duncan, & Desimone, 1993; Luck, Chelazzi, Hillyard, & Desimone, 1997; Treue & Maunsell, 1996). This does not emerge in recordings from striate visual cortex, where RFs are too small to contain multiple objects (Moran & Desimone, 1985).

Although influential to our understanding of human vision, evidence of biased competition from research with humans remains sparse and inconclusive. For example, results show that stimuli manipulations that should induce ambiguity – such as bringing a distractor close to an attended target – cause an increase in amplitude of the N2pc, an ERP index of spatial attention (Luck & Hillyard, 1994; Luck et al., 1997; Brisson, Robitaille, & Jolicoeur, 2007; Rashal et al., 2022; Luck & Ford, 1998; Eimer, 1996). This suggests that the N2pc reflects mechanisms recruited to resolve ambiguity, possibly silencing small-RF cells so that large-RF cells represent only the attended object (Luck et al., 1997). In line with this, the distractor positivity (P_d_), a sub-component of the N2pc, has been linked to direct suppression of distractor representation (Hickey, Di Lollo, & McDonald, 2009; Weaver, van Zoest, & Hickey, 2017). However, these findings are also consistent with other computational principles and other neurophysiological theories of attention, such as divisive normalization (Reynolds & Heeger, 2009), priority map theory (Bisley & Goldberg, 2010), or the free energy / predictive coding framework (Friston, 2009). To date, there has been no opportunity to concretely identify a relationship between selective attention and ambiguity resolution using EEG, for lack of a clear, independent measure of representational ambiguity.

This has motivated development of measures of representational ambiguity from human brain signals. Seminal fMRI work contrasted simultaneous vs. serial object presentation, assuming that competition should emerge only when stimuli are presented at the same time. Simultaneous presentation was found to produce less activity in extrastriate visual cortex, with attention reducing this difference, and this was interpreted as evidence of a bias on competition (Kastner, De Weerd, Desimone, & Ungerleider, 1998; Kastner & Ungerleider, 2001). However, this interpretation is problematic. First, when stimuli are presented concurrently, reductions in brain activity may be the product of adaptation, rather than competition, to maintain neural activity within a functional range (Reynolds & Heeger, 2009). Second, sequential presentation may allow for short-duration investment of resources into the processing of each sequential object, ultimately creating more brain activity over a fixed interval (Kupers, Kim, & Grill-Spector, 2024). Finally, sequential presentation may result in eye movements to bring attended objects closer to the fovea, decreasing the effect of visual crowding (Rosenholtz, 2024).

To overcome these issues, fMRI researchers have recently turned to multivariate pattern analysis (MVPA; Haxby, Connolly, & Guntupalli, 2014). Here, brain activity patterns for isolated vs. combined stimuli are compared, and results show that when two objects are viewed, the deployment of attention to one object shifts the pattern of brain activity toward that elicited by this object when it is presented alone (Reddy, Kanwisher, & Van Rullen, 2009; Doostani, Gholam-Ali Hossein-Zadeh, Cichy, & Vaziri-Pashkam, 2024; Cohen & Tong, 2015). This delivers a powerful index of the resolution of representational ambiguity. However, unlike EEG, fMRI does not provide a distinct, time-resolved index of selective attention, and observed changes in voxel patterns may reflect post-attentive cognitive operations like working memory, decision making, or response selection. A clear link between selective attention and ambiguity resolution has therefore not yet been demonstrated using fMRI, for lack of a clear, independent measure of selective attention.

There are additional unresolved issues in our understanding of biased competition. The theory has developed from studies employing animal electrophysiology, and specifically from observations of a small subset of visual cells whose RFs are sized just right—large enough to encompass two distinct stimuli, but not so large that they cover too much of the visual field and make baseline measurements impossible. This leaves its scope unclear. Does biased competition have a wider influence in the brain? Separately, there is debate regarding exactly how competition is resolved.

Does this rely entirely on distractor suppression (Luck et al., 1997), on target enhancement (Kastner, Pinsk, De Weerd, Desimone, & Ungerleider, 1999; Muller et al., 2006), or some combination (Hickey, Di Lollo, & McDonald, 2009)?

Here, we report results from 2 neuroimaging experiments that test core predictions from biased competition theory and characterize the neural mechanisms underlying ambiguity resolution. We do so by measuring the N2pc index of attention while concurrently assessing representational ambiguity using MVPA. We had 3 hypotheses. First, we tested the idea that ambiguity in the neural representation of visual objects recruits attention. Our specific expectation was that when a target and distractor are both viewed, a stronger attentive response – and therefore larger N2pc – will be required when separate, individual presentation of these objects generates similar brain activity patterns. Second, we tested the proposal that attention acts to resolve this kind of representational ambiguity. Here, our expectation was that N2pc amplitude should predict the shifts in brain activity patterns described above, reflecting a disambiguation of the target representation. Third, we expected ambiguity resolution to involve both target enhancement and distractor suppression. To test this, we identified how N2pc predicted change in the representation of both targets and distractors, linking effects to both target-elicited N2pc and distractor-elicited P_d_.

In testing these hypotheses, we introduce a new methodological approach to the analysis of multivariate neuroimaging data that we call *representational alignment modelling.* Experiment 1 employs alignment modelling of EEG, leveraging the temporal accuracy of this measure to identify the time-course of attention-mediated ambiguity resolution. Experiment 2 employs alignment modelling of concurrently-recorded EEG and MRI, allowing us to localize effects in brain space.

## Results

Participants in both experiments (n = 44, n = 31) completed two tasks in interleaved blocks. In the search task they reported the category of an object within a cued, coloured annulus at one of 4 locations, without moving their eyes, while ignoring non-targets including one recognizable distractor (Fig 1a; Fig S1a). In the benchmark task, they viewed sequential displays containing a single object and reported image repetition (Fig S1b). The same image set (E1: 160 images; E2: 120 images) was employed in both tasks, and the benchmark results were used in analysis of search results. In Experiment 1 we collected 128-channel EEG while participants completed these tasks, and in Experiment 2 we collected 64-channel EEG and concurrent fMRI.

**Figure 1.**
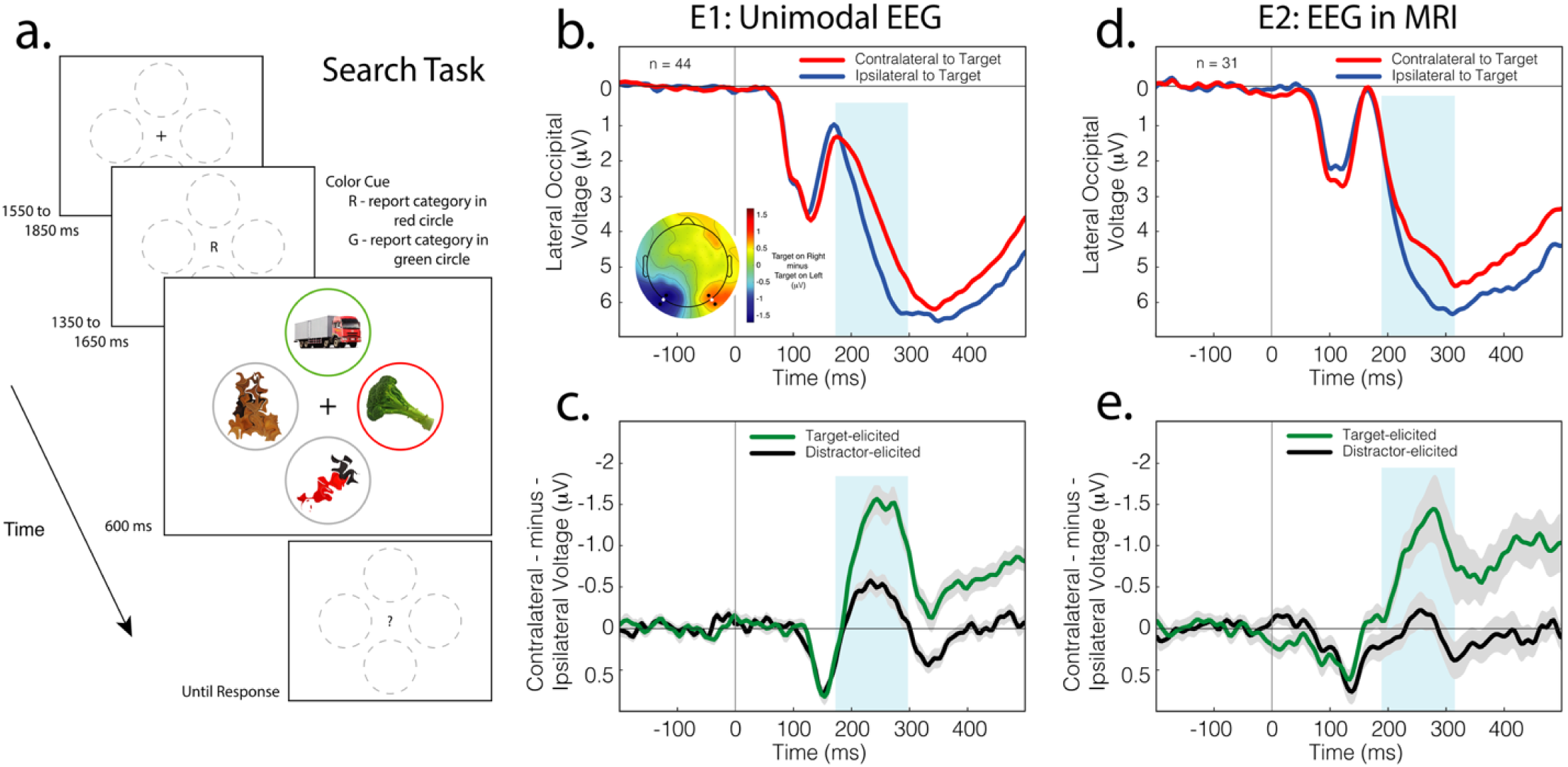
Experiment 1 task and ERP results. A.) Search task. The target was cued by a central letter, either ‘R’ or ‘G’ to denote red or green. Participants reported the category - tool, vehicle, animal, or food - of the object presented in the cued circle via right hand button response. Additional image examples and a schematic of the benchmark task are provided in Figure S1. B.) ERP results from Experiment 1. The N2pc is evident as a negativity in the posterior waveform recorded over cortex contralateral to an attended stimulus. The blue box identifies a 125 ms interval centred on the cross-conditional peak of the N2pc. The topographical plot reflects the voltage difference between conditions when the target was in the left versus right visual field; the N2pc expresses as a negativity in the left hemisphere but a positivity in the right hemisphere due to the subtraction. Contralateral and ipsilateral ERPs reflect mean signal across the 2 sets of 3 lateral and posterior electrodes identified in the topography by larger marker. C.) Ipsilateral-minus-contralateral difference waves from Experiment 1. The N2pc is evident as negative deflection beginning around 175 ms post-stimulus. Shading here and in other panels of this figure indicates bootstrapped SEM. D.) ERP results from Experiment 2. Results here reflect signal at the lateral posterior electrodes identified in white in the topography presented in panel A. E.) Ipsilateral-minus-contralateral difference waves from Experiment 2.

### Measuring attentional selection in the N2pc

In each trial of the search task, the target and distractor appeared either left or right of fixation or directly above or below it. When the target was presented laterally, the recognizable distractor was presented on the vertical, and vice versa (Fig 1a). This design was employed because stimuli on the vertical meridian of the visual field evoke equal activity in both cortical hemispheres, and do not produce the lateralized EEG voltage underlying N2pc and P_d_ (Woodman & Luck, 2003; Hickey, Di Lollo, & McDonald, 2009). This allowed us to isolate lateralized, attention-related EEG activity elicited solely by the target or recognisable distractor (Hickey, McDonald, & Theeuwes, 2006). We approached the data with the idea that, in each trial, correct deployment of attention to the target would generate a target-elicited N2pc and a distractor-elicited P_d_, reflecting target selection and distractor suppression respectively, but that misdeployment of attention to the distractor would generate a distractor-elicited N2pc. These possible outcomes are illustrated in Figure S2.

Averaging across trials, results show a target-elicited N2pc in both experiments (Fig 1b, 1d). A small distractor-elicited N2pc emerged in Experiment 1 (Fig 1c) but did not reach statistical significance in Experiment 2 (Fig 1e). In all cases, lateral voltage in the N2pc latency interval was preceded by earlier, positive-polarity lateral voltage, reflecting hemispheric sensory imbalance. In the search task, there was a difference in the complexity and spatial frequency of the complete object, presented to one side of the screen, and the unrecognisable, degraded object presented to the other (Fig 1a), and this generates short-latency laterality in the ERP (Pomerleau, Fortier-Gauthier, Corriveau, Dell’Acqua, & Jolicoeur, 2014). The insensitivity of this effect to the behavioural relevance of the evoking stimulus distinguishes it from N2pc and P_d_. We measured amplitude of the lateralized evoked response as the mean across a 125 ms window centred on the cross-conditional peak (E1: 179 – 298 ms; E2: 189 – 315 ms).

### Measuring the relationship of representational ambiguity and attentional selection using EEG

Our first hypothesis was that attentional mechanisms reflected in target-elicited N2pc are recruited when the target and distractor evoke similar neural responses in the benchmark task. To test this, we identified the similarity of mean benchmark EEG responses elicited by all possible pairs of the 160 objects in our stimuli set (Fig 2a). We generated 12,880 time-courses, each reflecting the electrode-wise correlation of evoked activity for a stimulus pair. We then associated each search trial with the corresponding correlation time-course based on target and distractor identity. We used mixed linear models to relate the trial-wise amplitude of N2pc to the correlation of corresponding benchmark responses.

**Figure 2.**
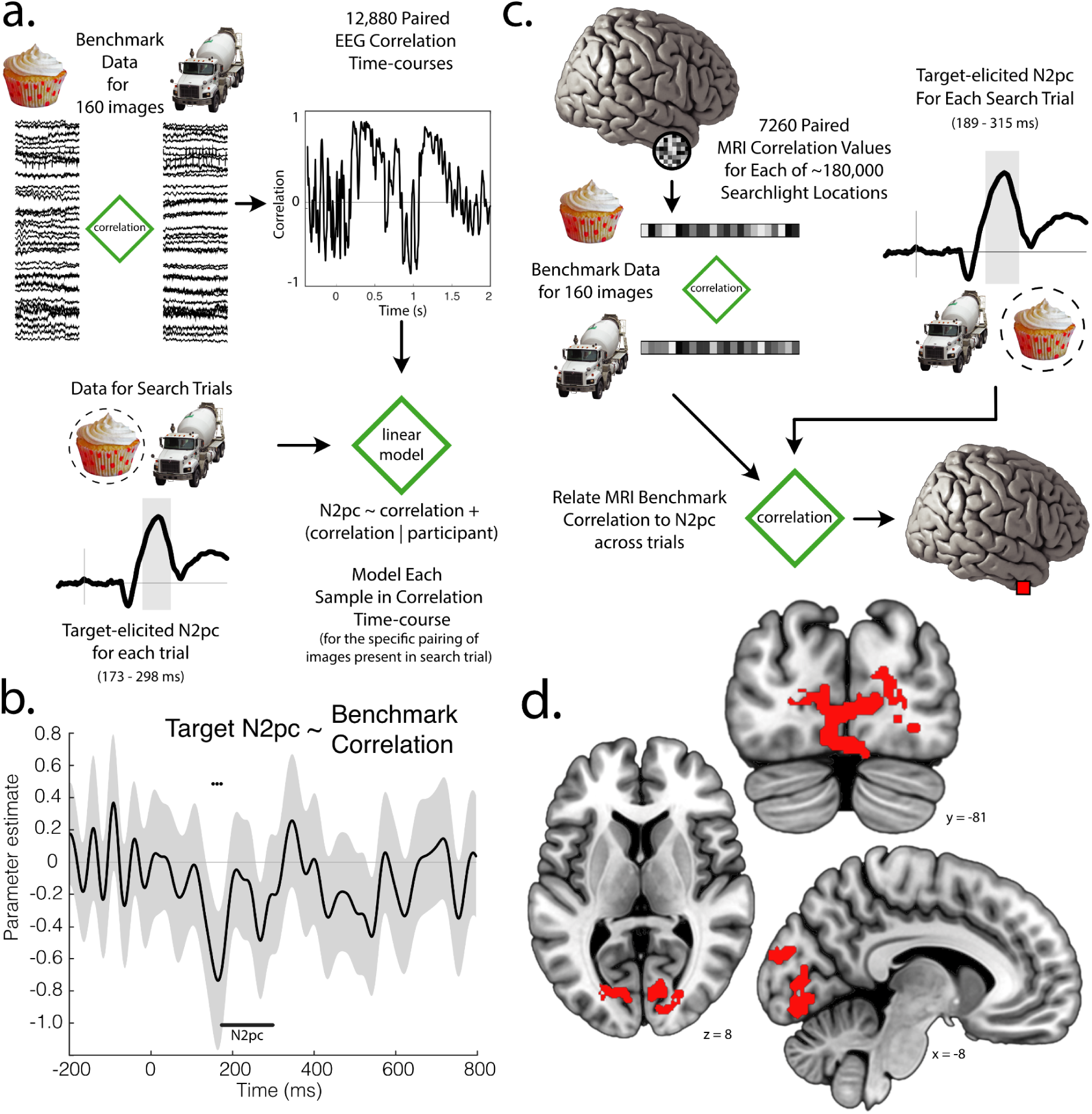
Linking representational ambiguity to N2pc. a.) Analysis schematic for Experiment 1. Benchmark EEG responses for all stimuli were correlated to create benchmark correlation time-courses. Each trial in the search task was associated with the benchmark correlation time-course corresponding to the pairing of target and distractor images. N2pc amplitude was modelled as a function of the benchmark correlation of target and distractor images. b.) The relationship between EEG benchmark correlations and target-elicited N2pc. The N2pc was measured as mean voltage at the fixed latency identified in the figure. Line shading reflects the uncorrected bootstrap 95% CI for that parameter estimate and FDR corrected statistical significance for each sample is denoted with circle markers above the line. For illustration, results are low-pass filtered at 30 Hz (5^th^ order Butterworth IIR). Corresponding analysis of distractor-elicited N2pc is illustrated in Figure S3 and garnered no significant results. c.) Analysis schematic for Experiment 2. Benchmark voxel activity was extracted from each searchlight for each object image and used to create 7260 parametric maps of benchmark correlation values. Each trial was associated with the benchmark correlation map corresponding to the pairing of target and distractor images. At each searchlight location, N2pc amplitude across trials was correlated with benchmark correlation values across trials. d.) The relationship between fMRI benchmark correlations and target-elicited N2pc. Greater benchmark correlation values predicted larger target-elicited N2pc in posterior visual cortex. Corresponding analysis of distractor-elicited N2pc garnered no results that survived correction for multiple comparisons. Details on voxel clusters emerging from this analysis are provided in Table S1.

As illustrated in Figure 2b, when the target appeared laterally in the search array, the correlation of target and distractor benchmark data at around 180 ms predicted the amplitude of the target-elicited N2pc. This supports the idea that selective attention, as reflected in N2pc, is recruited when target and distractor activate a similar population of cells, competing for representation in overlapping neural resources. This effect did not emerge in analysis of the distractor-elicited lateralized response (Fig S3).

### Measuring the relationship of representational ambiguity and attentional selection using concurrently recorded EEG and fMRI

Analysis of EEG/fMRI results from Experiment 2 localized brain structures where representational ambiguity triggers the recruitment of attention. The logic of analysis followed that above, but with benchmark correlations derived from searchlight analysis of fMRI (Kriegeskorte, Goebbel, & Bandettini, 2006). We began by generating whole-brain benchmark maps for each of the 120 object images employed in Experiment 2. We subsequently created a sphere around each voxel, extracted voxel-wise results in this sub-space, and cross-correlated these searchlight responses for all object pairs (Fig 2a). This yielded 7,260 benchmark correlations for each of ∼1.8 x 10^5^ searchlights. We then turned to results from the search task, using the GLMsingle toolbox (Prince et al., 2022) to estimate single-trial fMRI responses, and associated each search trial with the corresponding searchlight correlations based on target and distractor identity. For each participant, and for each searchlight, we Pearson-correlated the benchmark relationship with N2pc amplitude across trials, inserting the resulting values into new brain volumes at the location of the searchlight centre. Cluster-based permutation analysis was used to correct for multiple comparisons (Stelzer, Chen, & Turner, 2013) and group-level results appear in Figure 2b. These show that representational ambiguity in early visual areas predicts target-elicited N2pc amplitude. Areas showing this relationship included V1, V2, and V3, as well as ventro-occipital-temporal areas hV4, VO1, and VO2 (Wang, Mruczek, Arcaro & Kastner, 2015). Separate analysis of distractor-elicited results identified no areas surviving family-wise error correction.

### Tracking the attention-mediated resolution of representational ambiguity using representational alignment modelling of EEG

The results above establish that target-elicited N2pc is recruited when stimuli compete for representation in early visual cortex. This is consistent with the idea that N2pc acts to resolve ambiguity by biasing object representations (Desimone & Duncan, 1995; Luck et al., 1997), but does not demonstrate this principle, and the remainder of the paper is dedicated to testing this hypothesis.

We use a new analytic approach to identify the impact of N2pc on object representations, which we call representational alignment modelling. This tool extends existing fMRI methods (Reddy, Kanwisher, & Van Rullen, 2009; Doostani et al., 2024; Cohen & Tong, 2015; Kriegeskorte, Mur, & Bandettini, 2008) and builds from the 2-stage approach described above, where brain activity patterns from benchmark data inform analysis of search data. In the preceding sections we have used benchmark data to generate a single correlation value, which was employed as a scalar summary of representational ambiguity. In alignment modelling, analysis remains in multivariate space: benchmark patterns are used to directly predict activation patterns in search trials. The logic of the approach is illustrated in Figure 3. In analysis of EEG, each search trial generates a pattern of evoked voltage across electrodes. Benchmark data reflects the response to single images when they are attended and competition is low. Alignment modelling uses optimized mixed linear models to identify how the pattern evoked in search trials is predicted by the target and distractor benchmarks Unsurprisingly, search data strongly aligns with both target (Fig S4a) and distractor benchmark data (Fig S4b). This direct alignment is of limited interest, and emerges because both search and benchmark responses capture event-related activity evoked by similar stimuli. As illustrated in Figure S4, this occurs even when benchmark data is randomly selected and unrelated to the actual trial stimuli. This kind of control analysis – where search data is modelled with randomly selected benchmarks – is applied in all alignment modelling of EEG and denoted in figures by broken lines.

**Figure 3.**
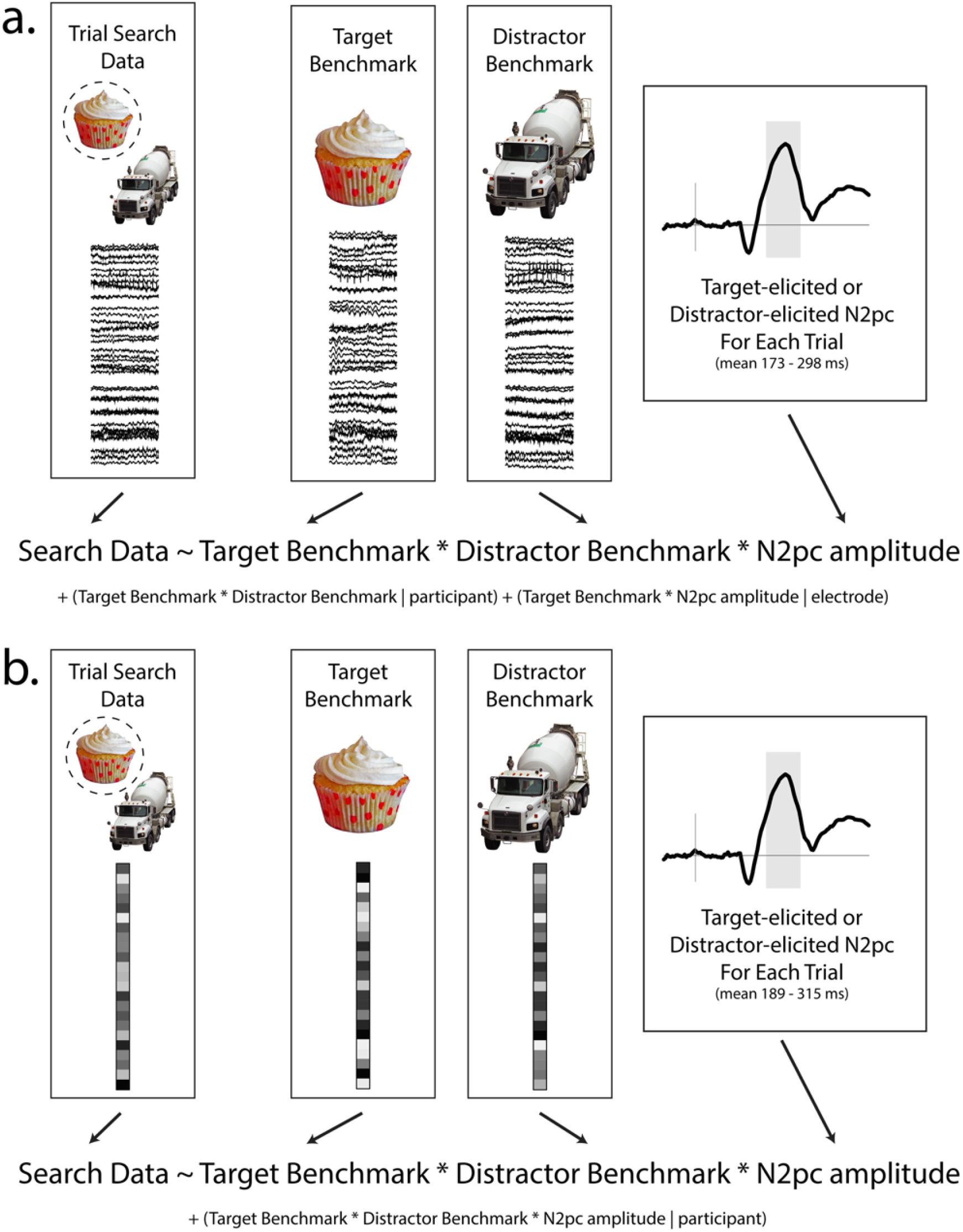
Logic of representational alignment modelling. a.) In Experiment 1, the voltage pattern at each latency of the evoked response for each search trial is modelled as a function of target benchmark, distractor benchmark, N2pc amplitude, and the interaction of these factors. B.) In Experiment 2, the voxel pattern in each searchlight location for each trial is modelled as a function of target benchmark, distractor benchmark, and N2pc amplitude, and the interaction of these factors.

#### The effect of target-elicited N2pc amplitude

The power of alignment modelling emerges when we consider the interaction of benchmark data with N2pc amplitude. As illustrated in Figure 4a, the parameter estimate for the interaction of target-elicited N2pc and target benchmark data shows a sharp negative deflection around 380 ms post-stimulus. This parameter has negative polarity because the N2pc itself has negative polarity, and it indicates that a larger target-elicited N2pc predicts closer alignment of the search results with the target benchmark. Simultaneously, a positive deflection in the parameter estimate for the interaction of target-elicited N2pc and distractor benchmark data emerges, conveying that a larger target-elicited N2pc predicts misalignment of the search results with the distractor benchmark. These effects are absent in control analyses, and they show that spatial attention resolves representational ambiguity by enhancing target representation and suppressing distractor representation.

**Figure 4.**
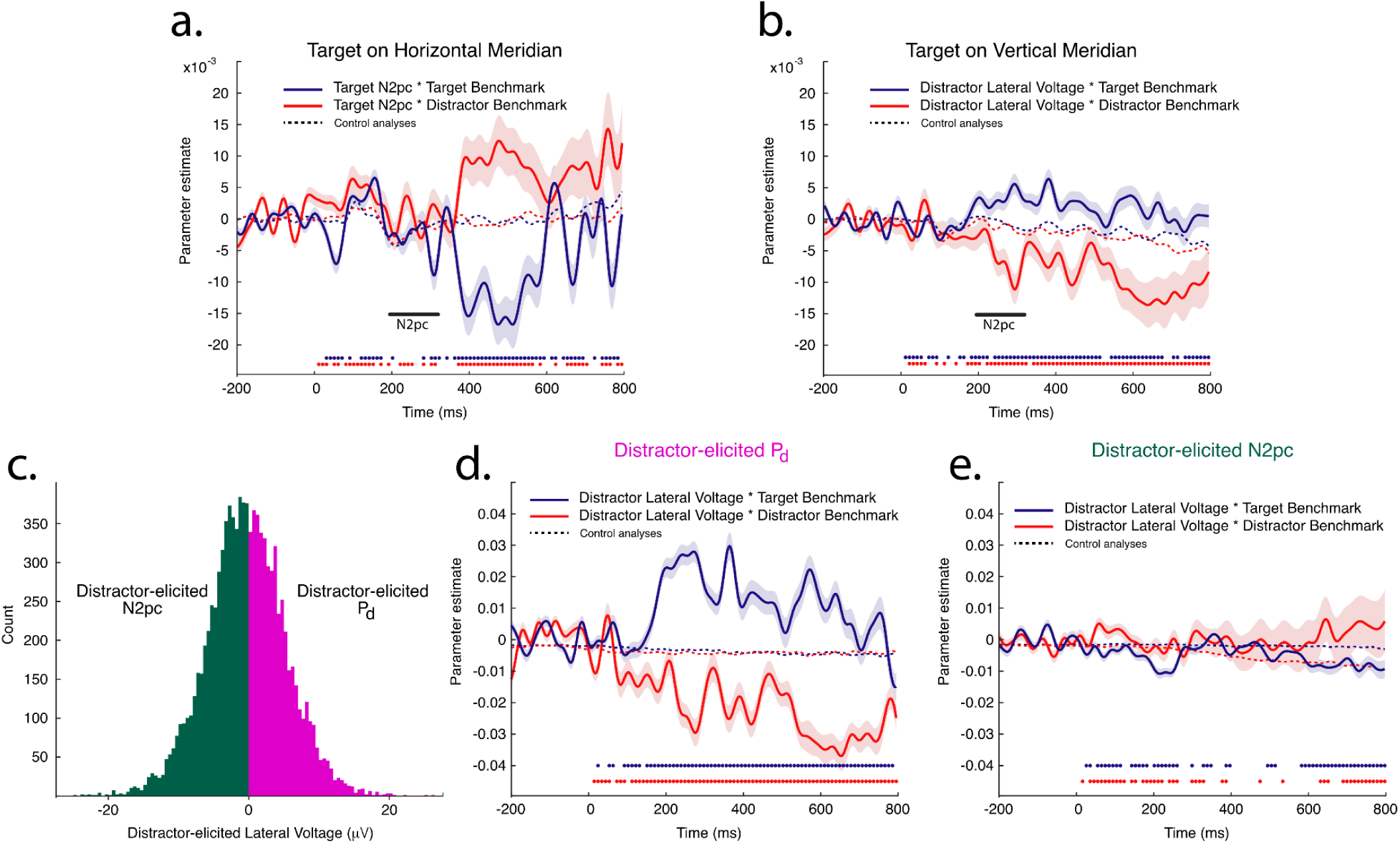
Using alignment modelling to track the resolution of representational ambiguity in EEG. In all panels, shading reflects the uncorrected 95% bootstrap CI for each parameter estimate. FDR-corrected significance is denoted in circle markers for each sample at the bottom of each panel. See methods for details on derivation of statistics. For illustration, parameter estimates are low-pass filtered at 30 Hz (5^th^ order Butterworth IIR). a.) Interaction of target-elicited N2pc and benchmarks when the target is presented laterally and evokes lateralized brain activity. Negative parameter polarity in this and the next panel indicate that lateral voltage amplitude predicts the alignment of benchmark data with search data. b.) Interaction of distractor-elicited N2pc and benchmarks when the distractor is presented laterally and evokes lateralized brain activity. c.) The distribution of lateral distractor-elicited voltage observed in trials collapsed across participants. In 46.8% of trials the lateral voltage had positive polarity, and in the remainder of trials the lateral voltage had negative polarity. d.) Alignment modelling of trials where the lateral response to the distractor had positive voltage. Greater positive voltage predicted alignment of search data with the target benchmark and misalignment with the distractor benchmark. e.) Alignment modelling of trials where the lateral response to the distractor had negative voltage.

#### The effect of distractor-elicited lateral brain activity: Distractor selection or suppression?

Our design allows us to measure lateralized EEG activity elicited during attentional processing of the distractor, and Figure 4b illustrates results from alignment modelling employing distractor-elicited lateral EEG signal. The pattern of results contrast with those for target-elicited N2pc: the interaction of distractor-elicited lateral activity and the target benchmark shows positive polarity, indicating misalignment of the search data with the target benchmark, whereas the interaction with distractor benchmark shows negative parameter polarity, indicating alignment of the search data with the distractor benchmark.

These results are consistent with two possibilities. It may be that as the distractor-elicited N2pc becomes increasingly negative, the search data increasingly expresses the distractor representation, reflecting the misallocation of attention to the distractor (Figure S2b). However, as illustrated in Figure 1c, the lateral response evoked by distractors was not robustly negative in polarity. In fact, as illustrated in Figure 4c, there were nearly as many trials where the distractor evoked positive lateral voltage as negative. As noted above, positive-polarity voltage over lateral occipital cortex is known as the P_d_ (Hickey, Di Lollo, & McDonald, 2009) and is linked to attentional suppression during target resolution (Weaver, van Zoest, & Hickey, 2017; Hilimire, Hickey, & Corballis, 2012; Gaspelin et al., 2023). This raises an alternative account for the pattern of results illustrated in Figure 4b: when attention is deployed to the target on the vertical meridian, ambiguity is resolved in part through suppression of the distractor representation, and this is reflected in amplitude of the distractor-elicited P_d_ (Figure S2c).

To distinguish between these interpretations, we separated trials into subsets based on whether the distractor elicited negative or positive lateral voltage and we modelled each subset using the predictors in Figure 3a. The mean number of distractor-lateral trials eliciting positive voltage was 107 (14.0 SD) per participant and the mean number of distractor-lateral trials eliciting negative voltage was 121 (16.7 SD). As illustrated in Figure 4d, when distractor-elicited P_d_ was observed, the amplitude of this signal predicted alignment of the search data with the target benchmark and misalignment of the search data with the distractor benchmark. This pattern did not emerge when negative voltage was observed (Fig 4e). This suggests that the distractor-elicited P_d_ tracks the resolution of representational ambiguity through inhibition of the distractor representation. The functional role of the distractor-elicited, negative-polarity signal is less clear, as we do not see evidence that this substantively improves representation of the distractor.

### Tracking the attention-mediated resolution of representational ambiguity using representational alignment modelling of concurrently recorded EEG and fMRI

#### How target-elicited N2pc resolves representational ambiguity

The alignment modelling of unimodal EEG described above identifies how complementary target-centered and distractor-centered mechanisms resolve representational ambiguity, and identifies the time-course of these effects. Concurrent EEG / fMRI results from Experiment 2 allow us to identify how these mechanisms impact representations in brain space. The logic of alignment modelling of EEG/MRI follows that described above, but benchmark and search patterns now come from searchlight fMRI (Kriegeskorte, Goebbel, & Bandettini, 2006). Model parameters for the interaction of N2pc with benchmark data are calculated for each searchlight and placed into a new brain volume at the searchlight centre. The primary product of this analysis is a set of parameter maps where each voxel reflects the statistical z-score for a single parameter derived from mixed linear modelling. We applied parametric threshold-free cluster enhancement (pTFCE; Spizák, Spizák, Zunhammer, Bingel, Smith, Nichols, & Kincses, 2019) to integrate evidence from the emergence of contiguous voxel clusters before applying false discovery rate (FDR; Benjamini & Hochberg, 1995) correction for multiple comparisons. Our analysis generated six parametric maps of interest. The first two appear in Figure 5 and show two effects: the ability of target-elicited N2pc to predict alignment of search data with the target benchmark, and the ability of target-elicited N2pc to predict misalignment with the distractor benchmark.

**Figure 5.**
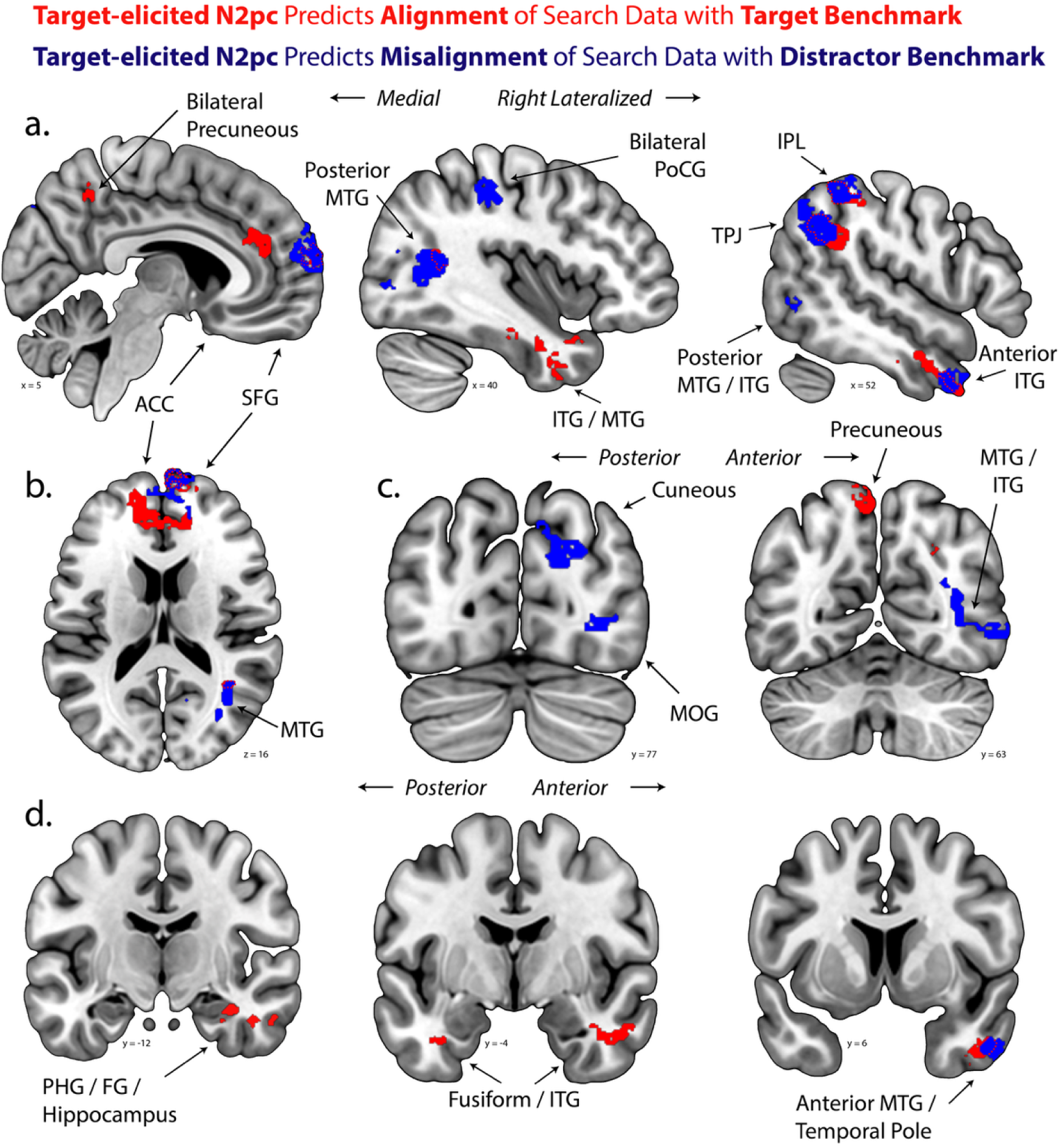
Using alignment modelling to track the resolution of representational ambiguity in combined EEG / fMRI. The interaction of target-elicited N2pc and the target benchmark is identified in red. The interaction of target-elicited N2pc and the distractor benchmark is identified in blue. Illustrated here are the positive relationship with target benchmark and the negative relationship with distractor benchmark. See Figure 7a for the negative relationship between target-elicited N2pc and the target benchmark; no positive relationship between target-elicited N2pc and the distractor benchmark emerged from analysis. a.) Coronal slices identifying effects on midline cortex extending into right temporal lobe. b.) Horizontal slice identifying effects in medial prefrontal cortex. c.) Coronal slices identifying effects in posterior occipital and temporal lobes. d.) Coronal slices identifying effects in anterior temporal lobe and hippocampus. Further information on clusters illustrated here is provided in Tables S2 and S3. ACC anterior cingulate gyrus; SFG superior frontal gyrus; MTG middle temporal gyrus; ITG inferior temporal gyrus; IPL inferior parietal lobe; TPJ temporo-parietal junction; MOG middle occipital gyrus; PHG parahippocampal gyrus; FG fusiform gyrus.

Before further considering these results, it is important to note that alignment modelling of EEG / fMRI is less sensitive to effects in early visual cortex with small-to-medium receptive fields than it is to effects in later, large-RF visual cortex and non-retinotopic cortex. This is because each object appears at each of 4 possible stimuli locations in our benchmark task, and these results are summarized before being used in modelling. The benchmarks therefore do not capture the precise spatial location of the object in retinotopic space. There are two key benefits to this: it means that lateralized, attention-related brain activity does not consistently contribute to the benchmark EEG patterns, and it means that the alignment of benchmark to search data cannot simply reflect the similarity of brain activity elicited by objects at the same location. However, it also means that the benchmark pattern does not robustly capture the response to the eliciting object in retinotopic cortex, as the mean response collapses across instances where this object was presented at different locations.

This is not to say that the N2pc has no effect on retinotopic cortex. We have shown that short-latency ambiguity in early visual cortex is associated with emergence of the N2pc, and the N2pc itself prominently reflects activity in retinotopic lateral occipital cortex (Fig 1b; Hopf et al., 2000). Our analytic technique, however, captures the effect of N2pc on representation in downstream cortical areas. Accordingly, results in Figure 5 largely emerge in anterior occipital, posterior parietal, temporal, and frontal areas.

Results from Figure 5 show that target-elicited N2pc predicts the alignment of search data with the target benchmark, and misalignment with the distractor benchmark, across a large and overlapping network. In Figures 5a and 5b we see that target-elicited N2pc predicts alignment of search data with the target benchmark in the right inferior parietal lobule, right temporal parietal junction, across bilateral temporal lobes, and in the bilateral superior frontal gyrus. Strikingly, target-elicited N2pc also predicts misalignment of search data with the distractor benchmark in these same areas. These complementary effects are not made necessary by the analysis – this is an emergent quality of the impact of attention on neural representation, not of the statistical approach - and Figures 5c and 5d show the emergence of clusters showing selective effects of target enhancement and distractor suppression.

#### How distractor-elicited EEG laterality resolves representational ambiguity

Corresponding analysis of distractor-elicited EEG laterality is illustrated in Figure 6. Experiment 1 identified that the lateral response to the distractor is a combination of trials with contralateral negativity and trials with contralateral positivity, and that only trials with contralateral positivity – or P_d_ – robustly predicted shifts in EEG activity patterns. As such, in Figure 6 we reference effects to the emergence of contralateral P_d_, showing how increasing P_d_ predicts the *alignment* of search data with target benchmark data and *misalignment* with distractor benchmark data. It is important to keep in mind that this labelling reflects interpretation, and that the relationship can alternatively be described as a function of contralateral negativity, showing how increasing *N2pc* predicts *misalignment* of search data with target benchmark data and *alignment* with distractor benchmark data.

**Figure 6.**
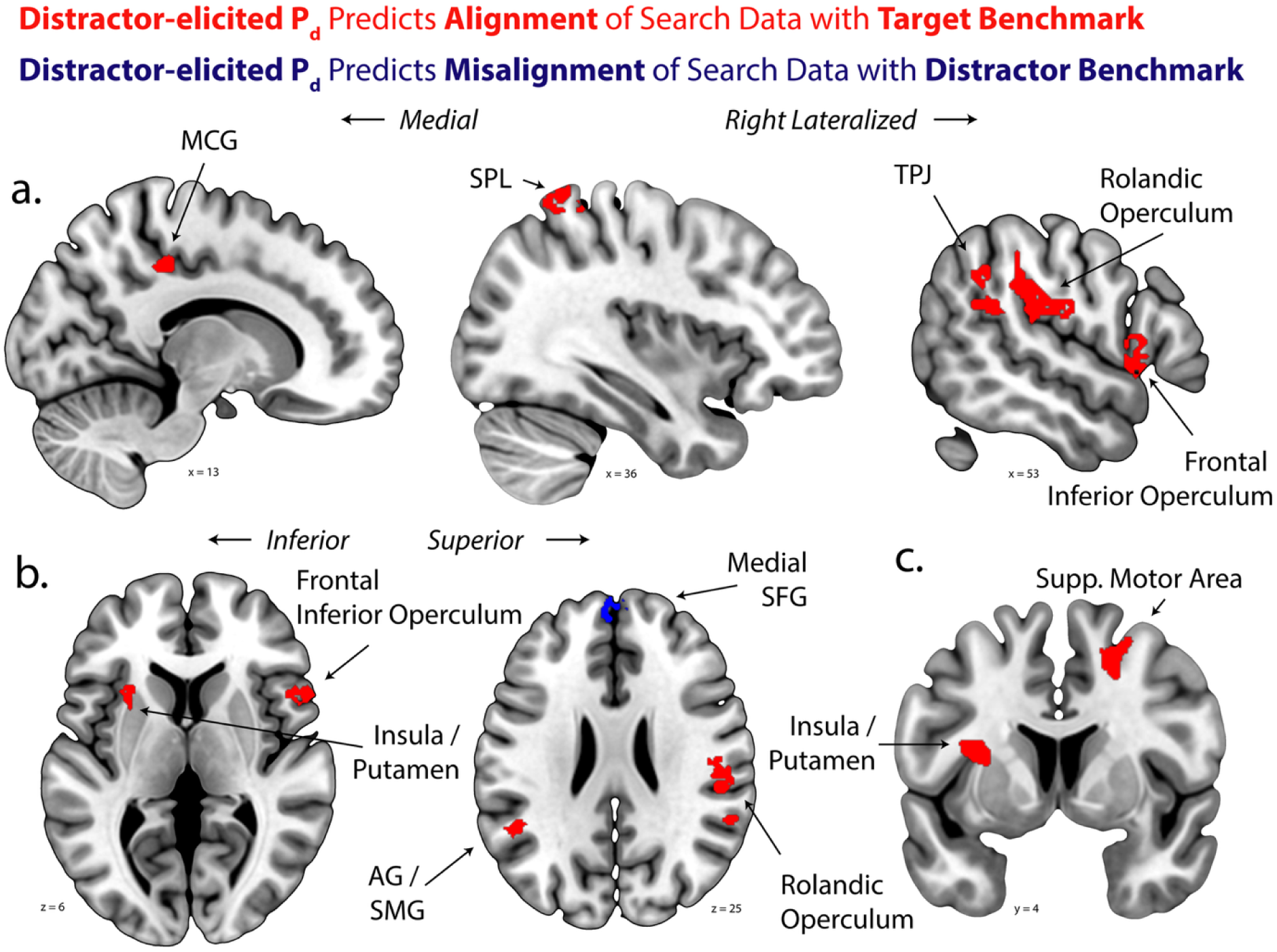
Using representational alignment modelling to track the impact of distractor processing on the resolution of representational ambiguity in combined EEG/fMRI. The interaction of distractor-elicited P_d_ and the target benchmark is identified in red. The interaction of distractor-elicited P_d_ and the distractor benchmark is identified in blue. Illustrated here are the positive relationship with the target benchmark and the negative relationship with the distractor benchmark. No negative relationship with the target benchmark emerged in analysis and the positive relationship with distractor benchmark appears in Figure 7b. a.) Coronal slices identifying effects on midline cortex extending into right temporal lobe. b.) Horizontal slice identifying effects in the bilateral opercula, medial prefrontal cortex, and left angular gyrus. c.) Coronal slices identifying effects in the insula and supplementary motor area. Further information on clusters illustrated here is provided in Tables S4 and S5. MCG middle cingulate gyrus; SPL superior parietal lobule; TPJ temporo-parietal junction; AG angular gyrus; SMG supramarginal gyrus; SFG superior frontal gyrus.

Two effects are observed: distractor-elicited P_d_ predicts alignment with the target benchmark and misalignment with the distractor benchmark. This emerges in a variety of brain areas, prominently including the right superior parietal lobule, right TPJ, and bilateral frontal operculum (Fig 6a).

Misalignment with the distractor benchmark emerges in a single bilateral cluster in the medial superior frontal gyrus. The propensity toward a broad relationship between distractor-elicited P_d_ and pattern alignment with the target benchmark is consistent with results from analysis of Experiment 1 illustrated in Figure 4b, where a stronger effect on alignment with the target benchmark is also observed.

#### When N2pc predicts a cost in representation

The results above track the benefit of selective attention through enhanced representation of attended objects and inhibition of unattended objects. However, N2pc also predicted counterintuitive negative effects on the representation of attended stimuli. The first of these effects is illustrated in Figure 7a and reflects the relationship between target-elicited N2pc and the misalignment of search data with the target benchmark. A single cluster emerges spanning the hippocampus and anterior lingual and fusiform gyri. One interpretation is that this reflects increasing use of mnemonic visual representations when participants do not rapidly and efficiently deploy attention during task completion. In our experimental design the target array is presented for only 600 ms. Analysis is limited to experimental trials that are correctly completed and we measure N2pc at a fixed latency. When participants do not deploy attention efficiently to the target, resulting in a low-amplitude target-elicited N2pc, but ultimately respond correctly, there may be greater need for a detailed mnemonic reproduction of the stimulus display. The hippocampus and proximal anterior medial visual areas are known to play a role in visualization and reactivation of episodic visual memories (Staresina, Henson, Kriegeskorte, & Alink, 2012), and this may explain the increased alignment of search data to target benchmark in these areas when the target-elicited N2pc is small.

**Figure 7.**
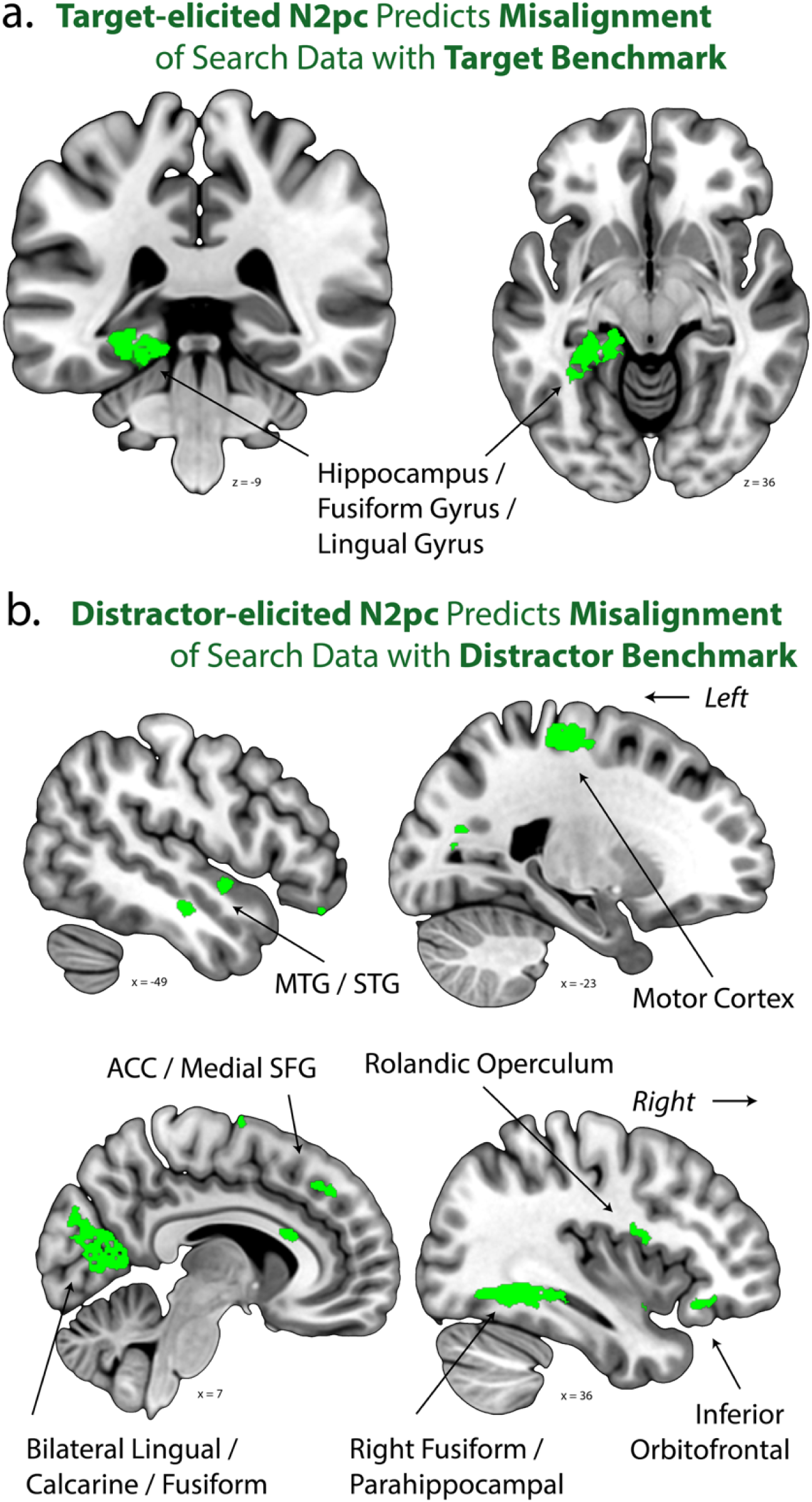
a.) Using alignment modelling to track the counter-intuitive impact of target-elicited N2pc on the misalignment of search data with the target benchmark. See main text for interpretation. Further information on voxel clusters is provided in Table S6. b.) Using alignment modelling to track the counter-intuitive impact of distractor-elicited N2pc on the misalignment of search data with the distractor benchmark. Note that this relationship could be alternatively labelled as the impact of distractor-elicited P_d_ on the alignment of search data with the distractor benchmark; the labelling employed in the figure reflects the interpretation offered in the main text. Further information on voxel clusters is provided in Table S7. MTG middle temporal gyrus; STG superior temporal gyrus.

The second of these effects is illustrated in Figure 7b and reflects the ability of distractor-elicited N2pc to predict misalignment of search data with the distractor benchmark. This is a clearer finding; when attention is initially misdeployed to the distractor, but participants are able to correctly respond to the target, results summated over the interval of BOLD fMRI show a reactive suppression of the distractor representation (Hickey & Peelen, 2015; Geng, 2014). Here, such suppression emerges in the left motor cortex, reflecting inhibition of the distractor-associated right-hand response; in the middle and anterior temporal lobe, reflecting suppression of the semantic and conceptual representation of the target; and across occipital cortex, reflecting suppression of early visual representation of the distractor.

## Discussion

This paper addresses a fundamental question in neuroscience: how does the brain resolve representational ambiguity created when visual objects compete for limited neural resources? We test three hypotheses: a.) that representational ambiguity recruits attention, b.) that attention resolves ambiguity by biasing competitive interactions between representations, and c.) that this involves complementary mechanisms acting on target and distractor representations. We find that when participants view two objects, and one is attended, the amplitude of N2pc is larger if the objects elicit similar activity patterns when presented individually in separate trials. This effect emerges around 180 ms post-stimulus, suggesting that the emergence of representational ambiguity early in the visual processing stream creates the need for attentional resolution. Our EEG/MRI results confirm this, showing that ambiguity in striate and extrastriate visual cortex predicts the target-elicited N2pc (Figure 2d; Luck et al., 1997).

To test how attention resolves ambiguity and the role of distractors in this process, we have developed a technique called *representational alignment modelling*. This allows us to identify how ERP indices of attentional mechanisms predict shifts in object representations instantiated in multivariate brain activity patterns. When participants viewed displays containing a target and distractor, larger target-elicited N2pc predicted that the activity pattern looked more like that elicited by the target alone, and less like that elicited by the distractor alone. These effects both emerge ∼400 ms after stimulus onset in a neural network that includes the IPL and angular gyrus / TPJ; anterior and posterior temporal cortex; and anterior cingulate and dorsomedial prefrontal cortex.

This set of brain areas overlaps with a network linked to semantic representation and control (Binder, Desai, Graves, & Conant, 2009). ‘Hub-and-spoke’ models propose the ATL binds semantic content from other regions (Lambon Ralph et al., 2016; Jackson, 2021). Sub-divisions of ATL have specialization in this context: the middle and inferior temporal lobes - which show an effect of attention in our results - are sensitive to semantic content conveyed by images rather than written or spoken words, whereas the superior temporal lobe - which does not show an effect of attention in our results - is responsive to abstract words (Hoffmann, Binney, & Ralph, 2015). We find that the effect of attention is stronger in the right ATL, consistent with hemispheric bias for image-based semantics (Neudorf, Gould, Mickleborough, Ekstrand, & Borowsky, 2022; Sevostianov et al., 2002).

Target-elicited N2pc also predicts representational changes in right TPJ / angular gyrus and posterior middle temporal gyrus, consistent with roles for these structures in the ventral attention network (VAN; Corbetta & Schulman, 2002) and in semantic manipulation (Humphreys & Lambon Ralph, 2015; Noonan, Jeffries, Visser, & Lambon Ralph, 2013). Additional effects in dorsomedial prefrontal cortex (dMPFC) and the dorsal anterior cingulate (dACC) suggest involvement of semantic (Jackson, 2021) and executive control systems (Clairis & Lopez-Persem, 2023). Distinct but linked theoretical accounts propose that dmPFC and dACC support conflict monitoring (Botvinick, Braver, Barch, Carter, & Cohen, 2001), calculation of the expected value of control (Shenhav, Botvinick, & Cohen, 2013), the monitoring of error likelihood (Brown & Braver, 2005), and the orchestration of cognitive exploitation versus exploration (Kolling, Behrens, Mars, & Rushworth, 2012). It thus appears that as the target is more strongly attended, information about the attended object emerges in brain areas involved in semantic representation and control, and in the implementation of conflict monitoring and executive regulation.

Importantly, target-elicited N2pc predicts effects on both target and distractor representation, and these effects emerge with the same time course in a largely-overlapping network. This supports the view that the N2pc reflects not only target enhancement (Eimer, 1996; Wolfe, 1994) - possibly via neural tuning or signal gain (Treue & Maunsell, 1996; McAdams & Maunsell, 1999) - but also that N2pc implements a particular type of distractor suppression. This kind of distractor suppression is theorized to shelter the representation of attended objects by limiting the ability of the distractors to interfere with the target representation (Luck et al., 1997; Hickey, Di Lollo, & McDonald, 2009). This mechanism could be implemented through interneurons physically close to cells responsible for representation of the target, explaining why the N2pc is contralateral to targets (Eimer, 1996) yet predicts the quality of distractor representation, and why effects on target and distractor representation in the current data emerge in overlapping brain areas.

Target sheltering differs from the direct distractor suppression indexed in P_d_, suggesting that three discrete attentional mechanisms are reflected in N2pc and P_d_: target enhancement, target sheltering, and distractor suppression. Where target enhancement and target sheltering are closely linked, direct distractor suppression appears distinct. The P_d_ predicts earlier effects (∼200 ms) than the N2pc (∼400 ms), consistent with other evidence of rapid distractor suppression in naturalistic vision of real-world objects (Hickey, Pollicino, Bertazolli, & Barbaro, 2019). Anatomically, both N2pc and P_d_ predict target enhancement and distractor suppression in the right TPJ, but unlike N2pc, P_d_ predicts target enhancement in the insula, the Rolandic and frontal opercula, the SPL / AG / SMG, and the MCG, and predicts distractor suppression in the medial prefrontal cortex.

As discussed above, the right TPJ is a prominent structure in both semantic control and in the VAN, and the opercula are commonly co-activated with TPJ and are also included in descriptions of the VAN (Corbetta, Patel, & Schulman, 2008; Cook, Im, & Giaschi, 2025). The opercula are also closely linked to the insula, and play a role in the insula-centered ‘salience network’ that is thought to orchestrate broad cognitive response to sudden events like the onset of stimuli (Seeley, Schatzberg, Glover, Kenna, 2007). Posterior cingulate cortex, including the MCG, is the third prominent brain area in the salience network. The cluster in SPL / AG / SMG is possibly a reflection of the dorsal attention network (DAN), which is closely involved in the implementation of strategic attentional control (Corbetta & Schulman, 2002). Finally, as noted above, medial prefrontal cortex is broadly associated with executive control. The results thus suggest that effective direct distractor suppression drives improved target representation in attentional control systems and stops representation of the distractor in a brain area involved in conflict monitoring and executive control.

It is important to note the relationship between these results and our earlier findings, where we related N2pc amplitude to abstract conceptual information in the brain (Acunzo, Grignolio, & Hickey, 2025). The studies differ both in terms of motivation and methodology. In our earlier work, we used machine learning to identify the brain signals common to varying examples of a category, showing that N2pc predicted the emergence of such abstracted category information the inferior parietal lobe, the insula, and ventromedial prefrontal cortex, areas that are proximal but not overlapping with current clusters. Together, results suggest that attention impacts both object-specific and abstracted categorical semantics in distinct networks.

Figure 8 visually summarizes results presented in the paper. Attention, as reflected in the N2pc component of the visual ERP, is recruited when representation in early visual cortex is ambiguous. Ambiguity is resolved through the action of linked, complementary mechanisms indexed in the N2pc and P_d_ that enhance representation of the target, shelter the target from distractor interference, and suppress representation of the distractor (Luck et al., 1997; Hickey, Di Lollo, & McDonald, 2009). The action of these mechanisms in lateral occipital cortex has a broad downstream impact on semantic and executive processing. When the target-elicited N2pc is larger, and the target is more effectively attended, target enhancement and target sheltering mechanisms drive the emergence of unambiguous target representation in brain structures involved in semantic representation and executive control. At the same time, when the distractor-elicited P_d_ is larger, and the distractor is more effectively suppressed, this drives a.) more robust emergence of target information in an independent network associated with attentional control and salience, and b.) weaker emergence of distractor information in an area associated with high-level executive control. The results demonstrate the critical role of ambiguity resolution in orchestrating cognitive operations throughout the brain.

**Figure 8.**
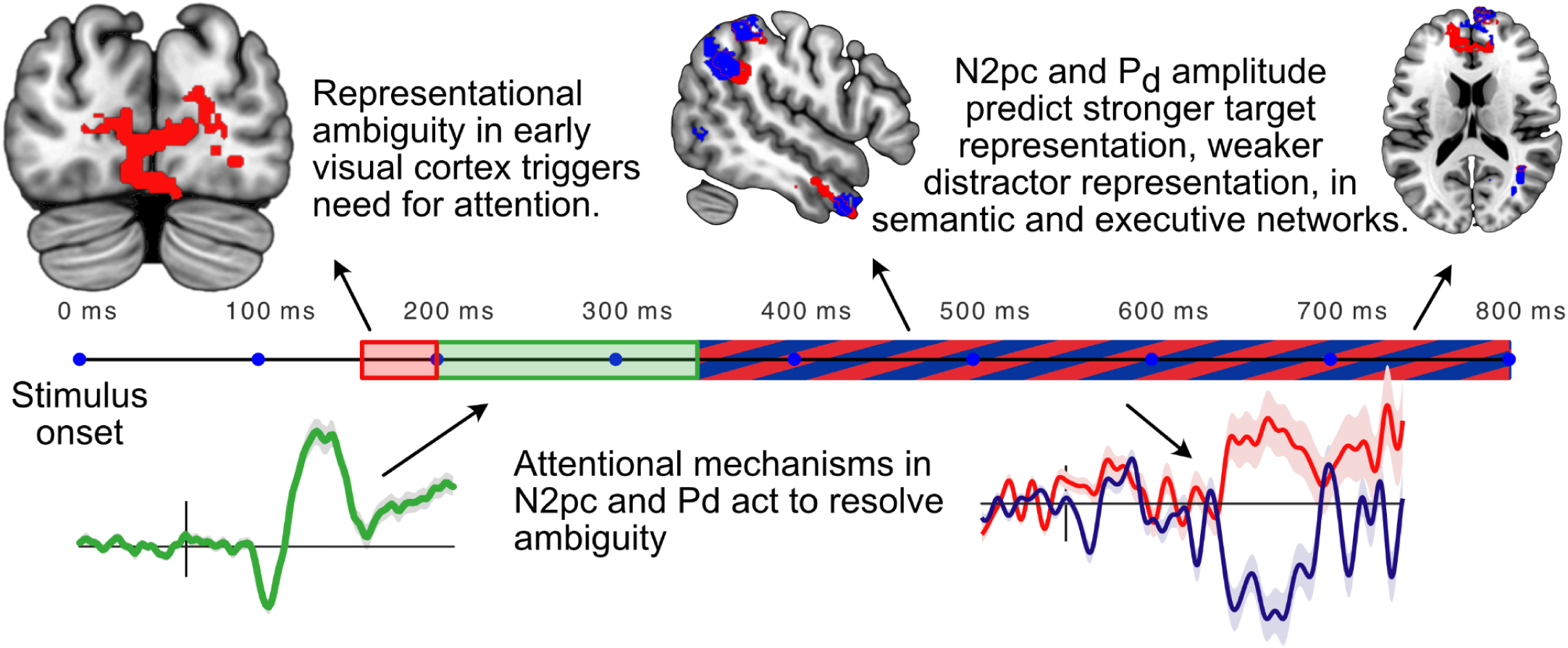
Summary of key results.

## Methods

### Human participants

All participants were recruited from the University of Birmingham community, provided written informed consent, reported normal or corrected-to-normal vision, and reported no history of neurological or psychiatric disorders. All demographics collected from experimental participants are reported and results can be expected to generalise to the European population from which the sample was taken. Experimental procedures were approved by the University of Birmingham STEM ethics committee and the Health and Safety committee of the Centre for Human Brain Health. The data underlying the results described in this paper have been employed to test other hypotheses in other published work (Acunzo, Grignolio, & Hickey, 2025; Hickey, Grignolio, Munasinghe, & Acunzo, 2024).

#### Experiment 1

Forty-nine participants were recruited. Two were excluded from analysis due to excessive eye movement artefacts in the EEG (rejection of >40% of trials), two were excluded as results showed no consistent evidence of a target-elicited N2pc in the search task (ie. across trials, the lateral target-elicited signal showed contralateral positive polarity more often than negative polarity; per-participant p < 0.05, binomial test), and a single participant was excluded due to low accuracy in search task performance (<70% accuracy; final sample of 44; age mean = 21.8, std = 2.7; 23 female; 3 left-handed). The experiment took approximately 2.5 hours, including EEG preparation, task training, breaks, and debrief, and participants were compensated at £10/hour.

#### Experiment 2

Thirty-nine participants were recruited. Two participants were excluded from analysis due to excessive eye movement artefacts in the EEG (rejection of >40% of trials), 4 participants were excluded due to incomplete data collection due to technical issues or equipment failure, and 2 participants were excluded due to low accuracy in task performance (<70% accuracy; final sample of 31; age mean=23.2, std=3.8; 19 female; 3 left-handed). The experiment took approximately 4 hours, including EEG preparation, MRI screening, task training, breaks, and debrief, and participants were compensated at £15/hour.

### Experimental task

Participants in both experiments performed a search task and a benchmark task in interleaved blocks (4 blocks of 128 search trials in both experiments; 4 blocks of 185 benchmark trials in Experiment 1 and 4 blocks of 156 benchmark trials in Experiment 2; block order counterbalanced across individuals). In each experimental trial of the search task, they reported the semantic category of a target that appeared in a cued location and ignored a distractor that appeared in an un-cued location (Fig 1a; Fig S1). Each trial began with a fixation display on white background (1550 – 1850 ms, randomly selected from a uniform distribution) that contained 4 grey circles (E1: 5.7° visual angle; E2: 4.2°) directly above and below (E1: 7.4° visual angle; E2: 5.2°) a central fixation mark. The fixation mark was replaced by the letter ‘R’ or ‘G’ (1350 – 1650 ms), which indicated the red or green colour of the circle identifying the target in the subsequent search array. In the search array, images of complete objects appeared within the two circles that had red or green colour. The objects were tools, vehicles, animals, or foods, and were selected from an image set containing equal numbers of each category (E1: 160 objects in total; E2: 120 objects; Fig. S1a) taken from the BOSS database (Brodeur, Dionne-Dostie, Montreuil, & Lepage, 2010) or open online image repositories. Target examples were selected from each category randomly without replacement until the set was exhausted and the process reset. The other two circles in the display contained unrecognisable morphed versions of the same image set (Stojanoski & Cusack, 2014; max distortion 160, nsteps 4, iteration 9), selected in a similar manner. The search array sustained for 600 ms and the next trial began either immediately after response or after 1800 ms.

The experimental task was to identify which object category appeared in the cued circle with right hand button response. Two of the categories were associated with index finger response and the others with middle finger response, with the mapping of category to response counterbalanced across participants. In Experiment 1 response was made via a standard keyboard; in Experiment 2 response was via an MRI-compatible button box. The distractor was randomly selected from a category associated with response opposite that of the target.

In the benchmark task, participants viewed sequential displays that contained a single instance of a complete, recognisable object and 3 instances of unrecognisable morphed objects, each presented in a grey circle (Figure S1b). Each display was presented for 600 ms and was followed by a display containing only the 4 grey circles (500 – 1000 ms, uniform distribution). Object images were taken from the same stimulus set employed in the search task; the locations and sizes of the objects and circles were also as in the search task. Participants were asked to identify via right hand index finger response when the recognisable image was repeated across displays.

The experiment began with verbal instructions and participants subsequently completed training until they achieved ∼80% accuracy in the search task, which rarely took longer than a single block of trials. In Experiment 2 training took place outside of the scanner. In the search task, each experimental block was composed of 128 trials in random order, with each trial reflecting one combination of cue colour, target and distractor category, and target and distractor position. In Experiment 1, stimuli were presented via a 59.5 cm x 33.5 cm LCD monitor at 60 Hz and a distance of 60 cm. In Experiment 2, stimuli were presented via a ProPiXX projector at 100 Hz viewed via a mirror mounted on the scanner head coil. The experimental procedure relied on the PsychToolbox 3 toolbox for Matlab (Brainard, 1997).

### Data acquisition and preprocessing: Experiment 1

Electrophysiological data was acquired from sintered Ag/AgCl electrodes at 1 kHz using a Biosemi ActiveTwo amplifier. EEG was collected from 128 electrodes fitted in an elastic cap at equidistant encephalic sites, horizontal electrooculogram (HEOG) was collected from 2 electrodes located 1 cm lateral to the external canthi of the left and right eye, vertical electrooculogram (VEOG) was collected from 2 electrodes immediately above and below the right eye, and unused reference signals were collected from 2 electrodes placed over the left and right mastoid processes. EEG was resampled offline to 200 Hz, digitally filtered with symmetric Hamming windowed finite-impulse response kernels (high-pass at 0.05 Hz, −6 dB at 0.025 Hz; low-pass at 45 Hz, −6 dB at 50.6 Hz), referenced to the average of encephalic channels, and baselined on the 200 ms interval preceding stimulus onset. Noisy channels were visually identified and interpolated using spherical spline interpolation.

Independent component analysis (ICA; Hyvärinen & Oja, 2000) of combined EEG and EOG data was used to identify data variance resulting from eye movements. Trials with eye movements in the 500 ms interval following stimulus onset were removed from analysis and the ICA components reflecting eye artefacts were subsequently removed from the data.

### Data acquisition and preprocessing: Experiment 2

#### MRI

Whole-brain scanning employed a 3T Siemens Prisma MRI scanner and 64 channel head-coil. Functional data was acquired using an echo planar imaging (EPI) sequence with 57 axial slices of 84 x 84 voxels, field-of-view (FOV) 210 mm x 210 mm, slice thickness 2.5 mm, slice gap 0 mm, repetition time (TR) 1.5 s, echo time (TE) 35 ms, flip angle (FA) 71°, multi-band acceleration factor 3. Structural data was acquired using a T1-weighted MPRAGE sequence with 208 axial slices of 257 x 257 voxels, FOV 256 mm x 256 mm, slice thickness 1 mm, slice gap 0 mm, TR 2 s, TE 2.03 ms, FA 8°. For correction of other images, a field map was acquired using a double-echo sequence with 36 axial slices of 64 x 64 voxels, FOV 192 mm x 192 mm, slice thickness 3.75 mm, slice gap 0 mm, TR 400 ms, TE_1_ 4.92 ms, TE_2_ 7.38 ms, FA 45°. Four-lead vectorcardiogram (VCG) was recorded in the scanner from electrodes in a standard chest montage. VCG was recorded at MRI clock speed (5 kHz) and was synchronised with this signal.

MRI images were corrected for head movement, field map inhomogeneity, and slice-acquisition delay (using the middle slice as reference) before being normalised to Montreal Neurological Institute (MNI) space, interpolated to 2mm isotropic voxel size, and spatially smoothed using an isotropic Gaussian kernel (6mm full width half maximum). General linear model analysis (GLM) of the fMRI time-series data relied on the GLMsingle toolbox (Prince et al., 2022). GLMsingle identifies hemodynamic kernel functions, derives nuisance regressors, and chooses an optimal ridge regularisation shrinkage fraction for each voxel to produce an estimate of single-trial voxel-wise beta values. To align EPI volumes with the onset of visual stimulation, the EPI time series was resampled to 6.7 Hz (150 ms per image) using shape-preserving piecewise cubic interpolation (as implemented in the tseriesinterp.m function included in knkutils toolbox; https://github.com/cvnlab/knkutils). Analysis was constrained to voxels outside of the brainstem, the ventricles, and the cerebellum.

#### EEG

Electrophysiological data was acquired in the scanner from sintered Ag/AgCl electrodes using a Brain Products BrainAmp MR amplifier. EEG was collected from 63 electrodes fitted in a BrainCap MRI-compatible elastic cap at extended 10/20 encephalic sites and single-lead electrocardiogram (ECG) was recorded from an electrode located on the back 10 cm caudal to the top of the shoulder and 5 cm to the left of the spine. All data acquisition was at MRI clock speed (5 kHz) and synchronized with this signal. Preprocessing began with correction for MR gradient artifacts using BrainVision Analyzer 2 (v2.2). Gradient triggers sent by the scanner were used as time markers for artifact trains and continuous artifacts were corrected using template subtraction based on a sliding average of 21 TR intervals (with baseline correction). The EEG was subsequently resampled to 200 Hz and corrected for cardioballistic (CB) artefacts. The CB correction algorithm semi-automatically detects peaks from heartbeat signals. In the 124 experimental blocks analysed (4 for each of 31 participants), the signal quality and availability of heartbeat signals varied: VCG was used for correction in 52 blocks, ECG was used in 48 blocks, and the remaining 24 blocks relied on a heart-beat ICA component derived from the EEG. A running average of 21 beats was used to generate a template and the artifact time delay was automatically estimated using a 40 s interval. Subsequent digital filtering, baselining, noisy channel interpolation, ICA, and trial rejection parameters were as described for Experiment 1. In data from the search task, the average number of interpolated channels was 0.29 and the number of independent components removed was 2.0. The mean number of correct trial epochs containing excessive noise or eye movements was 27 (SD = 26), and the mean number of incorrect trials was 52 trials (SD = 36), leaving around 433 trials per participant for analysis (SD = 41).

### Quantification and analysis

#### Correlation of N2pc amplitude to benchmark similarity

In both experiments, analysis of search data depended on the identification of activity patterns derived from the benchmark task. Our initial analyses identified how voxel-wise similarity of the target and distractor predicted N2pc amplitude. This began with calculation of benchmark responses for each object image. Each object in our stimuli set appeared repeatedly in the benchmark task (∼4 times in Experiment 1; ∼5 times in Experiment 2), with each appearance at a different stimulus location (until all locations had occurred, when the process reset). We mean averaged trial voltage (Experiment 1) or single-trial beta weights within searchlights (Experiment 2; diameter = 9 voxels; constrained when bound by the edge of brain space, mean volume = 225 voxels) across repetitions of the same object image to create benchmark patterns. We then Pearson correlated all pairwise patterns at each latency or searchlight location to create benchmark correlation time-courses (Experiment 1) or benchmark correlation parameter maps (Experiment 2). We associated these benchmark correlations to each search trial based on the identity of the target and distractor, subsequently relating these values to N2pc amplitude across trials using mixed linear modelling (Experiment 1) or Pearson correlation (Experiment 2).

Statistical inference in these analyses employed resampling. In analysis of Experiment 1, we began by generating uncorrected 95% confidence intervals (CIs) for each parameter at each sample latency by randomly resampling the data with replacement 2500 times, modelling each dataset in each iteration using the predictor structure illustrated in Figure 2a. To meet the assumption of exchangeability, resampling was conducted such that idiosyncratic data clustering within model random effects was retained. We calculated statistical significance as the cumulative probability that the empirical parameter distribution for each sample would be equal to or greater than zero. These sample p-values were then FDR corrected for multiple comparisons with a critical alpha of 0.025 to implement a 2-tailed test (Benjamini & Hochberg, 1995).

In the corresponding analysis of Experiment 2, we calculated the cross-trial correlation between N2pc amplitude and benchmark correlation for each searchlight within each participant to generate per-participant parametric maps. We used a Montecarlo cluster permutation approach to assess the statistical significance of voxel clusters that emerged from this analysis (Oosterhof, Connolly, & Haxby, 2016; Maris & Oostenveld 2007). This relied on a cluster-defining threshold of p < 0.05 and a cluster statistic that reflected summed parameter value within each cluster (‘maxsum’).

#### Alignment modelling and model optimization

In alignment modelling, the pattern of activity observed in each trial of the search task was modelled as a function of the pattern of activity observed in the target and distractor benchmarks and the amplitude of the N2pc. In Experiment 1, alignment modelling was independently conducted for each 5 ms EEG sample. In Experiment 2, it was independently conducted for each fMRI searchlight.

The fixed effects in our models were defined by our experimental hypotheses but the random effect structure was empirically optimized. To this end, we first calculated the maximal model (Barr, Levy, Scheepers, & Tily, 2013) before iteratively simplifying the model using Akaike information criterion (AIC) as a measure of model improvement. For analysis of target-lateral trials in Experiment 1, the maximal model is illustrated in Eq. 1.

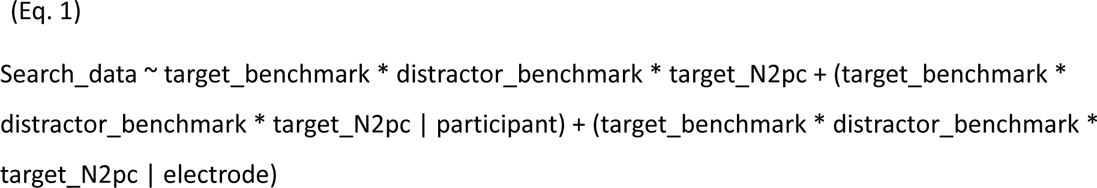

To illustrate how optimization occurred, our first simplification was to reduce the random effect for ‘electrode’ by removing the model term for interaction with ‘target_N2pc’, which reduced AIC. We then removed the additive term for ‘target_N2pc’, finding this increased AIC. We accordingly retained this factor and moved on within the random effect for ‘electrode’ to consider ‘distractor_benchmark’. We proceeded in this fashion to ultimately identify the model presented in Figure 3a. We repeated this process for samples at three latencies – 200, 300, and 400 ms post-stimulus – and found that optimization led to the same outcome in all instances.

We approached model optimization in Experiment 2 using similar logic. The maximal model was as presented in Eq. 1 except for the absence of a random effect for ‘electrode’. We calculated models for data in 10 bilaterally symmetric, equidistant searchlights spanning from early visual cortex anteriorly to prefrontal cortex. Our plan was to accept optimization steps when they improved model performance in 6 or more of these sample voxels. However, in practice, the best model at all sample locations was consistently the maximal model, which is illustrated in Figure 3b.

#### Statistical assessment

##### Experiment 1

Statistical inference from alignment modelling of EEG began with calculation of 95% CIs for the model parameters. For each temporal sample, we resampled the data with replacement 250 times, calculating the model illustrated in Figure 3a in each instance. Resampling was conducted such that data clustering within model random effects was retained.

We then conducted a control analysis to estimate the null result. For each iteration of this control analysis, we replaced the target and distractor benchmark patterns – which corresponded to benchmark data for the actual target and distractor stimuli present in the search trial – with target and distractor benchmarks generated by other randomly-selected objects in the stimulus set. We modelled this data with the predictor structure illustrated in Figure 3a, iterating this process 100 times for each temporal sample. Finally, we mean-averaged the set of resulting model parameters to generate the results illustrated by broken lines in Figure 4 and S4.

Statistical significance was calculated through comparison of the control results to the 95% CI of the model parameters. For each temporal sample, we calculated the probability that values equal to or exceeding the value generated in control analysis would be observed, given the empirical null parameter distribution generated via resampling. These sample p-values were then FDR corrected for multiple comparisons with a critical alpha of 0.025 to implement a 2-tailed test (Benjamini & Hochberg, 1995).

##### Experiment 2

The primary product of alignment modelling of combined EEG/fMRI results was a set of parameter map containing z-scores for each model parameter at each searchlight location. To integrate evidence provided by the emergence of spatial clusters, we applied probabilistic threshold free cluster enhancement (pTFCE) to these z-score parameter maps (Spisák, Spisák, Zunhammer, Bingel, Smith, Nichols, & Kincses, 2019) before using FDR (Benjamini & Hochberg, 1995) to correct for multiple comparisons within each map.

Analysis was largely conducted in Matlab (Mathworks, USA) using the EEGLAB (v.2024.1; Delorme & Makeig, 2004), SPM (v.25.01; Friston et al., 2007), and COSMOMVPA toolboxes (v. 2024; Oosterhof, Connolly, & Haxby, 2016). Mixed linear modelling of concurrent MRI/EEG results was conducted using the MixedModels.jl package (Bates, Alday, & Kokandakar, 2025) for the Julia programming language (Bezanson, Edelman, Karpinski, & Shar, 2017). Application of pTFCE relied on R (R core team, 2017) scripts provided by Spisák et al. (2019).

## Supporting information

Supplemental figures, results, and tables

## Acknowledgments

Our thanks to Vinura Munasinghe, Holly Ahmed, and Tia Cainer for support with EEG data collection; to Steve Mayhew for technical expertise on the recording of concurrent EEG/MRI; to Alicia Rybicki, Oscar Ferrante, and Wieske van Zoest for discussion; and to the University of Birmingham BlueBEAR high performance computing service (www.birmingham.ac.uk/bear) for computational resources.

## Author Credit

CH: Conceptualization, Software, Formal Analysis, Visualization, Writing – Original Draft, Writing – Review and Editing, Supervision, Project administration, Funding acquisition. DA: Investigation, Software, Formal Analysis, Visualization, Writing – Review and Editing. DG: Investigation, Formal Analysis, Writing – Review and Editing.

