## Supplemental figures, results, and tables for "Complementary attentional mechanisms for the resolution of representational ambiguity in the human brain"

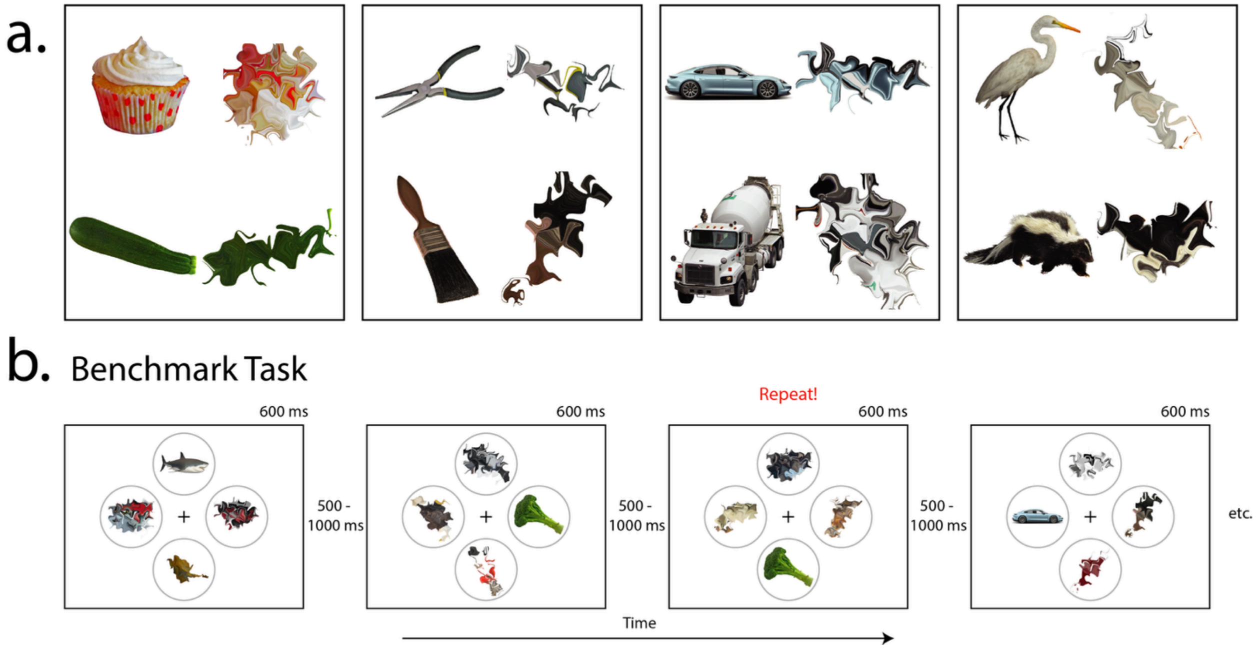


Figure S1, supporting Figure 1 – Experimental task and stimuli. a.) Examples of complete, recognizable objects and morphed, unrecognisable versions of the same images. In Experiment 1, the stimuli set was composed of 40 complete images in each of the 4 categories. In Experiment 2, a subset of 30 complete images per category was employed. b.) Benchmark task procedure. Each display contained a single recognisable object and three unrecognizable objects. Participants monitored the ongoing stream for repetition of the same specific image, regardless of location.

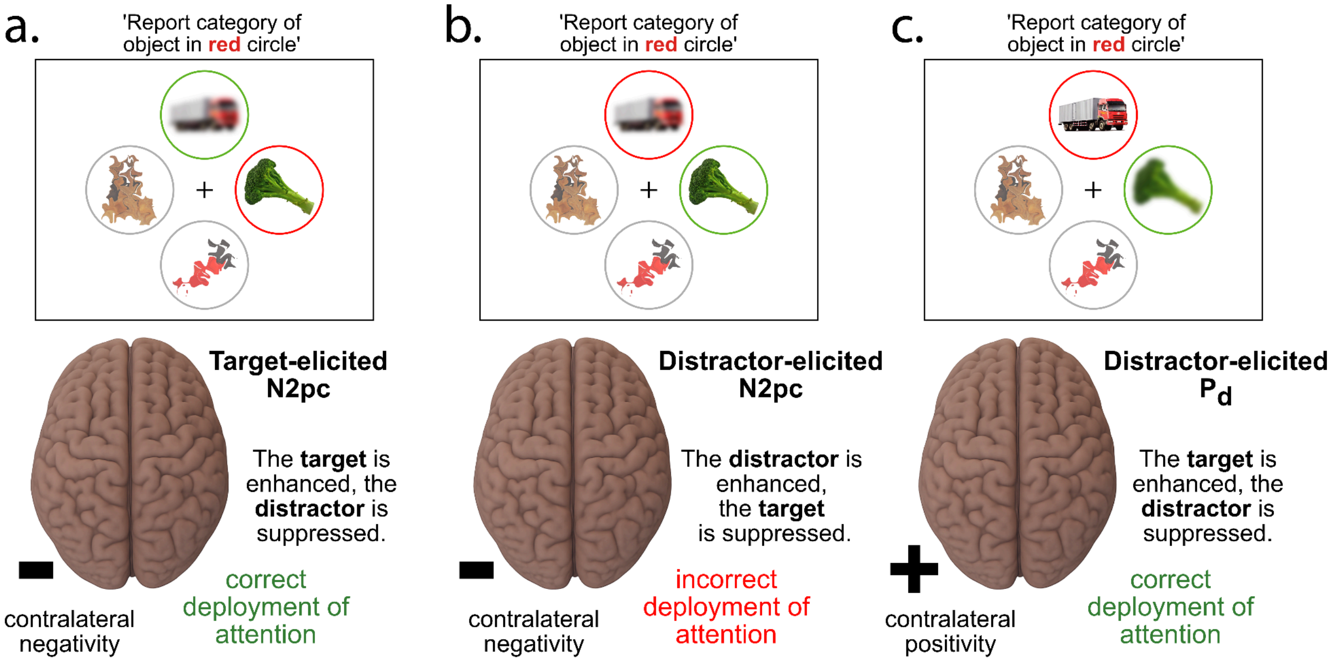


Figure S2, supporting Figure 1 – Possible outcomes in lateralized brain activity during search. a.) When the target appears at a lateral location in the visual field, its correct attentional selection will generate a target-elicited N2pc. Suppression of the distractor on the vertical meridian will not generate lateralized brain activity in the ERP. b.) When the target appears on the vertical meridian of the visual field, its correct attentional selection will not generate lateralized brain activity. Suppression of the distractor at a lateral location, however, will generate the distractor-elicited $P_d$. c.) When the distractor appears at a lateral location in the visual field, its erroneous attentional selection will generate a distractor-elicited N2pc. Suppression of the target on the vertical meridian will not generate lateralized brain activity.

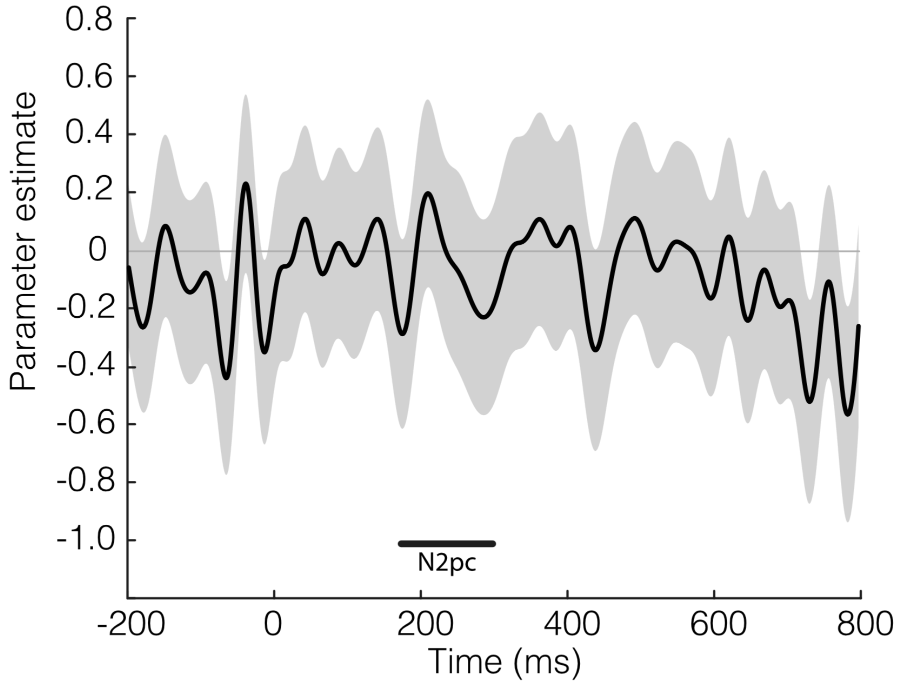


Figure S3, supporting Figure 2 – Additional results from modelling of N2pc amplitude as a function of EEG benchmark correlation timecourses. The relationship between benchmark correlation and distractor-elicited N2pc is illustrated. No effects survived correction for multiple comparisons.

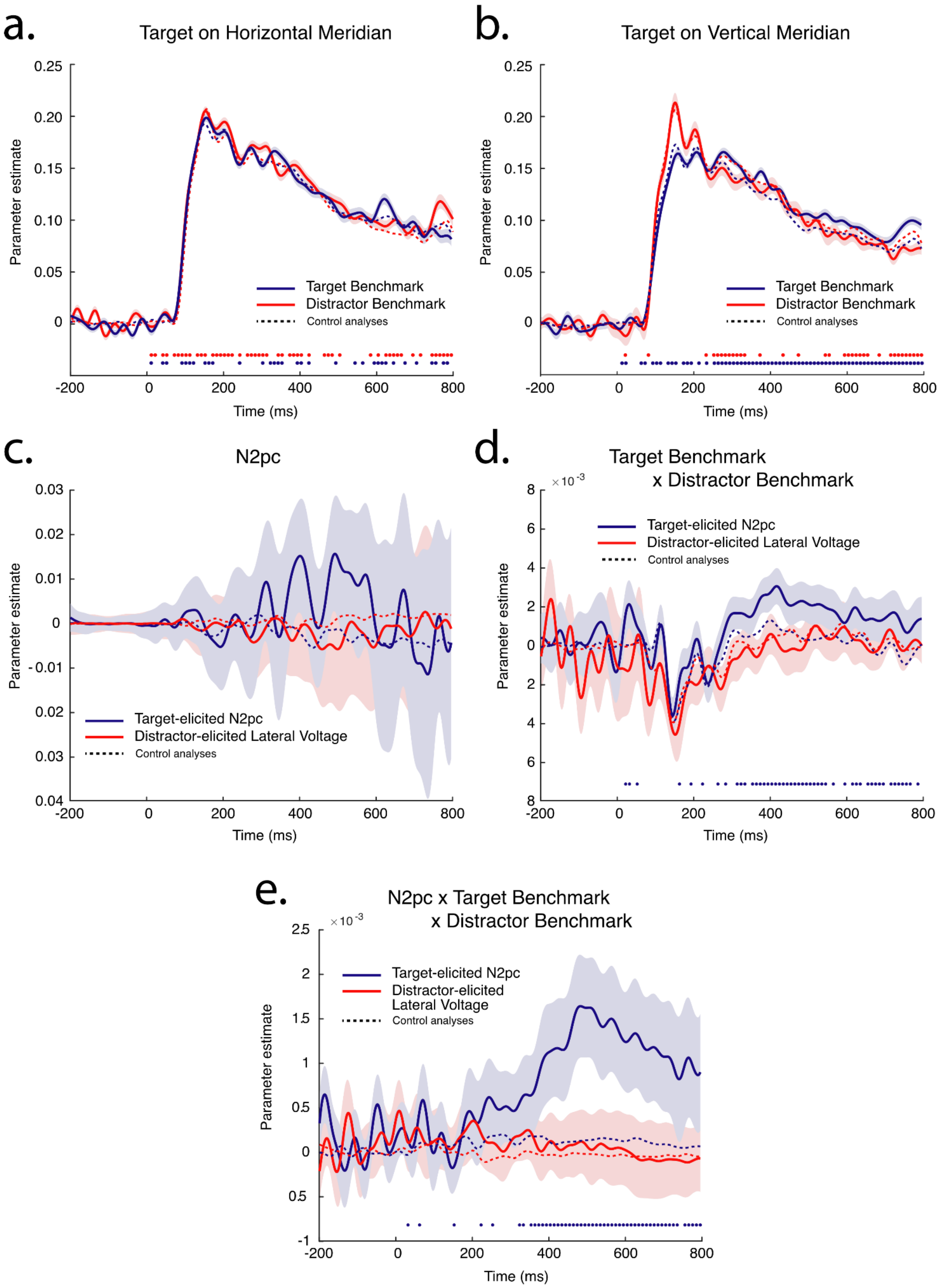

a.
Target on Horizontal Meridian
Parameter estimate
0.25
0.20
0.15
0.10
0.05
0
Target Benchmark
Distractor Benchmark
Control analyses
-200
0
200
400
600
800
Time (ms)
b.
Target on Vertical Meridian
Parameter estimate
0.25
0.20
0.15
0.10
0.05
0
Target Benchmark
Distractor Benchmark
Control analyses
-200
0
200
400
600
800
Time (ms)
c.
N2pc
Parameter estimate
0.03
0.02
0.01
0
- 0.01
0.02
0.03
0.04
Target-elicited N2pc
Distractor-elicited Lateral Voltage
Control analyses
-200
0
200
400
600
800
Time (ms)
d.
Target Benchmark
x Distractor Benchmark
× 10 -3
Parameter estimate
8
6
4
2
0
2
4
6
8
Target-elicited N2pc
Distractor-elicited Lateral Voltage
Control analyses
-200
0
200
400
600
800
Time (ms)
e.
N2pc x Target Benchmark
x Distractor Benchmark
× 10 -3
Parameter estimate
2.5
2
1.5
1
0.5
0
0.5
-1
Target-elicited N2pc
Distractor-elicited
Lateral Voltage
Control analyses
-200
0
200
400
600
800
Time (ms)

(previous page) Figure S4, supporting Figure 4 – Additional results from alignment modelling of EEG results in Experiment 1. Shading in all panels reflects parameter 95% CI and control analyses reflect mean of repeated analysis with benchmark data selected randomly, as described in the MS body. Samples where a parameter statistically differs from control analysis are identified with small circles near the bottom of the panel. Circles have color corresponding to each parameter; where no circles appear, no significant effect emerged. For illustration, parameter estimates are low-pass filtered at 30 Hz (5th order Butterworth IIR). a.) Direct alignment of search data with target and distractor benchmarks when the target is presented laterally and evokes the N2pc. Search data aligns closely with benchmark data because both reflect the evoked response to visual onset. b.) Direct alignment of search data with target and distractor benchmarks when the distractor is presented laterally and evokes lateralized brain activity. Note the change in organization from preceding panels; line color now reflects whether the target appeared at a lateral location (generating a target-elicited N2pc) or the distractor appeared at a lateral location (generating distractor-elicited lateral voltage) c.) Parameter estimates for the effect of N2pc amplitude on search data. d.) Parameter estimates for the effect of the interaction of target and distractor benchmarks on search data. e.) Parameter estimates for the effect of the threeway interaction of N2pc, target benchmark, and distractor benchmark on search data.

### Supplementary Results to complement Figure S3

Representational alignment modelling of EEG using the model structure illustrated in Figure 3a was conducted separately for trials where the target appeared at locations on the horizontal meridian (target-elicited N2pc), and trials where the target appeared at locations on the vertical meridian (distractor-elicited N2pc). For each of these analyses 7 parameter time-courses were generated, reflecting the model factors: target benchmark, distractor benchmark, N2pc amplitude, target benchmark * distractor benchmark, target benchmark * N2pc amplitude, distractor benchmark * N2pc amplitude, and the 3-way interaction.

Key results from the interaction of target-elicited and distractor-elicited N2pc with target and distractor benchmark patterns are illustrated in Figure 4c and 4d. Additional, less critical results from alignment modelling are illustrated in Figure S3.

Results from the direct alignment of the target and distractor benchmark are illustrated in Figures S3a and S3b. As noted in the main body of the paper, the search data aligns closely with the benchmark data as both reflect the evoked response to similar stimuli arrays.

Figure S3c illustrates the ability of N2pc amplitude to directly predict consistent pattern emergence in the search data. This is a sanity check; our modelling is based on the assumption that N2pc amplitude should have no power in predicting the emergence of any specific pattern in the search results. It is therefore reassuring to find that no effect emerges.

Figure S3d illustrates how the interaction of the target and distractor benchmarks predicts the search data. Results show a significant interaction, though only when the target appeared on the horizontal meridian of the display. This effect means that when the target benchmark was strongly reflected in the search data, so was the distractor benchmark, and we believe it reflects the influence of motivation and task engagement.

Figure S3e illustrates how the interaction of target benchmark and distractor benchmark are further modulated by N2pc amplitude. Results again show a significant effect, though only when the target-elicited N2pc is analyzed. In interpreting this effect, it is important to keep in mind the negative polarity of the N2pc. The pattern suggests that as the N2pc became smaller - and was characterized by more positive voltage - the interaction of target and distractor benchmarks described above became larger.

This effect is more easily understood if described as a product of increasing N2pc amplitude: when the N2pc became larger, and voltage was more negative, the interaction of target benchmark and distractor benchmark had less influence on the search results. When the N2pc is large, the target is well represented and the distractor is poorly represented, breaking the link between target and distractor reflected in the 2-way interaction.

| **Volume** (cubic mm) | **Location** (centroid; MNI XYZ) | **Peak Structure** | **Other Structures** |
|---|---|---|---|
| 10232 | 4 -84 8 | Right Calcarine | Calcarine_L Lingual_L Occipital_Sup_R Cuneus_L Cuneus_R Lingual_R Occipital_Sup_L Cerebelum_Crus1_R Vermis_6 Cerebelum_6_R Cerebelum_6_L Occipital_Mid_R |
| 472 | 26 -86 -4 | Right Lingual | Occipital_Inf_R Fusiform_R |

Table S1, supporting Fig. 2 – Voxel clusters where the similarity of target and distractor benchmark data predicts N2pc amplitude. Here and in subsequent supplementary tables only clusters of >= 24 cubic mm are listed. Structure correspondence is according to the automated anatomical labelling atlas version 3 (Rolls, Huang, Lin, Feng, & Joliot, 2020).

| **Volume** (cubic mm) | **Location** (centroid; MNI XYZ) | **Peak Structure** | **Other Structures** |
|---|---|---|---|
| 5160 | -4 46 20 | Left Anterior Cingulate | Cingulum_Ant_R Frontal_Sup_L Frontal_Sup_Medial_L |
| 2184 | 54 -48 34 | Right Supramarginal | Angular_R Parietal_Inf_R |
| 2136 | 48 2 -32 | Right Inferior Temporal | Temporal_Mid_R Temporal_Pole_Mid_R |
| 1128 | 8 66 16 | Right Medial Superior Frontal | Frontal_Sup_R Frontal_Sup_Medial_L |
| 1040 | 54 -40 52 | Right Inferior Parietal | SupraMarginal_R Parietal_Sup_R |
| 840 | 2 -52 48 | Left Precuneus | |
| 840 | -6 -64 62 | Left Precuneus | Precuneus_R |
| 608 | 42 -14 -24 | Right Inferior Temporal | Hippocampus_R Fusiform_R ParaHippocampal_R |
| 288 | 40 -54 16 | Right Middle Temporal | |
| 272 | 44 14 -28 | Right Superior Temporal Pole | Temporal_Pole_Mid_R |
| 216 | -36 -6 -32 | Left Fusiform | Temporal_Inf_L |
| 88 | 46 -18 -30 | Right Inferior Temporal | Fusiform_R |
| 88 | 40 4 -42 | Right Inferior Temporal | |
| 80 | 28 -62 36 | Right Superior Occipital | |

Table S2, supporting Fig. 6 – Voxel clusters where target-elicited N2pc amplitude predicts alignment of the search data with the target benchmark.

| **Volume** (cubic mm) | **Location** (centroid; MNI XYZ) | **Peak Structure** | **Other Structures** |
|---|---|---|---|
| 4648 | 50 -52 42 | Right Angular | Parietal_Inf_R SupraMarginal_R Parietal_Sup_R |
| 2672 | 6 64 18 | Right Medial Superior Frontal | Frontal_Sup_Medial_L Cingulum_Ant_R Frontal_Sup_R |
| 2120 | 20 -76 30 | Right Cuneus | Occipital_Sup_R Occipital_Mid_R |
| 1752 | 40 -30 46 | Right Postcentral | SupraMarginal_R Parietal_Inf_R |
| 1424 | 40 -58 12 | | Temporal_Mid_R(42) Occipital_Mid_R(2) |
| 848 | 50 10 -36 | Right Inferior Temporal | Temporal_Pole_Mid_R |
| 600 | 56 -66 0 | Right Inferior Temporal | Temporal_Mid_R |
| 448 | 32 -76 0 | Right Fusiform | Occipital_Mid_R Occipital_Inf_R Lingual_R |
| 408 | -32 -34 42 | | Postcentral_L Parietal_Inf_L |
| 64 | 14 70 22 | Right Superior Frontal | Frontal_Sup_Medial_R |
| 32 | 44 8 -38 | Right Middle Temporal Pole | Temporal_Pole_Mid_R Temporal_Inf_R(50) |

Table S3, supporting Fig. 6 – Voxel clusters where target-elicited N2pc amplitude predicts misalignment of the search data with the distractor benchmark.

| **Volume** (cubic mm) | **Location** (centroid; MNI XYZ) | **Peak Structure** | **Other Structures** |
|---|---|---|---|
| 3040 | 54×-48×32 | Right Supramarginal | Angular_R Temporal_Sup_R Temporal_Mid_R Parietal_Inf_R |
| 2656 | 52×-28×28 | Right Rolandic Operculum | SupraMarginal_R Rolandic_Oper_R Temporal_Sup_R |
| 1584 | -32×8×12 | Left Putamen | Insula_L Rolandic_Oper_L Precentral_L Frontal_Inf_Oper_L |
| 1320 | 22×2×52 | | Frontal_Sup_R Frontal_Mid_R Supp_Motor_Area_R |
| 1232 | 56×14×2 | Right Inferior Frontal, Opercular | Temporal_Pole_Sup_R Rolandic_Oper_R |
| 672 | 34×-52×66 | Right Superior Parietal | Postcentral_R |
| 608 | -48×-52×30 | Left Supranarginal | Angular_L Temporal_Mid_L Parietal_Inf_L |
| 352 | 14×-34×44 | Right Middle Cingulate | Precuneus_R |
| 216 | -2×-48×60 | Right Precuneus | Precuneus_L |
| 128 | 42×-44×64 | Right Superior Parietal | Postcentral_R |
| 112 | -6×60×34 | Left Medial Superior Frontal | |
| 56 | -54×-16×-8 | Left Middle Temporal | |
| 48 | 36×-44×62 | Right Superior Parietal | Parietal_Sup_R Postcentral_R |
| 24 | -28×14×-4 | Left Putamen | |

Table S4, supporting Fig. 7 – Voxel clusters where distractor-elicited N2pc amplitude predicts misalignment of the search data with the target benchmark.

| **Volume** (cubic mm) | **Location** (centroid; MNI XYZ) | **Peak Structure** | **Other Structures** |
|---|---|---|---|
| 1072 | -2×58×30 | Left Medial Frontal Superior | Frontal_Sup_Medial_R |
| 88 | 64×-58×-8 | | Temporal_Inf_R |

Table S5, supporting Fig. 7 – Voxel clusters where distractor-elicited N2pc amplitude predicts alignment of the search data with the distractor benchmark.

| **Volume** (cubic mm) | **Location** (centroid; MNI XYZ) | **Peak Structure** | **Other Structures** |
|---|---|---|---|
| 2224 | -24 -38 -10 | Left Fusiform | ParaHippocampal_L Cerebelum_4_5_L Hippocampus_L Lingual_L |
| 120 | -40 -60 10 | | Temporal_Mid_L Occipital_Mid_L |

Table S6, supporting Fig. 8a – Voxel clusters where target-elicited N2pc amplitude predicts misalignment of the search data with the target benchmark.

| **Volume** (cubic mm) | **Location** (centroid; MNI XYZ) | **Peak Structure** | **Other Structures** |
|---|---|---|---|
| 8884 | -1 -66 10 | Left Lingual | Calcarine_R Calcarine_L Fusiform_R Lingual_R Cuneus_L Cuneus_R(8) Occipital_Sup_R Occipital_Inf_R |
| 1045 | -23 -27 62 | Left Precentral | Precentral_L Postcentral_L Paracentral_Lobule_L |
| 267 | 4 21 18 | Bilateral Cingulate | Cingulum_Ant_R Cingulum_Ant_L |
| 105 | -50 -16 -18 | Left Middle Temporal | Temporal_Inf_L |
| 45 | -50 0 8 | Left Superior Temporal | |
| 42 | 10 -2 72 | Right Supplemental Motor Area | |

Table S7, supporting Fig. 8b – Voxel clusters where distractor-elicited N2pc amplitude predicts misalignment of the search data with the distractor benchmark.
